# Gene expression profiling of dendritic cell tolerance dysfunction in women with Systemic lupus erythematosus

**DOI:** 10.64898/2025.12.21.695630

**Authors:** Ana Laura Hernández-Ledesma, Evelia Lorena Coss-Navarrete, Sofia Salazar-Magaña, Diego Ramírez-Espinosa, Lizbet Tinajero-Nieto, Estefania Torres-Valdez, Angélica Peña-Ayala, Guillermo Félix-Rodriguez, Gabriel Frontana-Vázquez, Jair Santiago García Sotelo, Gosia Trynka, Florencia Rosetti, Selene Lizbeth Fernandez-Valverde, María Gutiérrez-Arcelus, Deshiré Alpízar-Rodríguez, Alejandra Medina-Rivera

## Abstract

Dendritic cells (DCs) are central regulators of immune tolerance, and disturbances in their phenotype and function contribute to the breakdown of self-tolerance in systemic lupus erythematosus (SLE). Tolerogenic DCs (tolDCs), which suppress autoreactive responses and promote peripheral tolerance, are a promising therapeutic focus in autoimmune diseases. Here, we analyzed the transcriptional profiles of *in vitro* generated DCs derived from monocytes of individuals with SLE and healthy controls to identify disease-specific disruptions in tolerance associated pathways. Interferon stimulated genes (ISGs) emerged as dominant markers across all cellular contexts, with monocytes exhibiting the most substantial enrichment; key ISGs (I*FI27, IFI44L, USP18, IFI6*) acted as central hubs in regulatory networks, underscoring their diagnostic and pathogenic significance. In tolDCs from SLE donors, lipid metabolism pathways were selectively altered, suggesting impaired synthesis of pro-resolving lipid mediators. Additionally, diminished *IL10RA* expression and dysregulated *IRF4* activity in SLE moDCs indicated intrinsic defects in IL-10 mediated tolerogenic differentiation. Together, these findings suggest that interferon driven transcriptional rewiring, impaired IL-10 signaling, and aberrant lipid metabolic programming converge to compromise DCs tolerogenic capacity in SLE. This highlights key mechanistic pathways that could be targeted to restore immune tolerance and reduce chronic inflammation.

## Introduction

Systemic Lupus Erythematosus (SLE) is an autoimmune disease characterized by immune dysregulation and increased production of autoantibodies targeting several tissues and organs, leading to multisystem involvement and a wide range of clinical manifestations (1). Affecting over 3.4 million people worldwide, SLE disproportionately impacts women of reproductive age (20-45 years) and people with Asian, African, and Hispanic/Latino ancestries (2–5).

Although the precise etiology of SLE remains unclear, its development results from the interplay of genetic, epigenetic, environmental, and hormonal factors (1,6). This interplay leads to the breakdown of immune tolerance, a key mechanism that triggers atypical and hyperreactive immune responses against self-antigens (7). In healthy individuals, immune tolerance is a dynamic process that prevents immune response against self-antigens, involving well-described central and peripheral mechanisms in both primary and secondary lymphoid organs. These include inactivation and control of self-reactive T and B cells through cell mechanisms such as apoptosis (8,9).

Dendritic cells (DCs) bridge innate and adaptive immune responses, and they can initiate either inflammatory or tolerogenic responses, depending on environmental stimuli (10,11). DCs are highly heterogeneous, with multiple subsets: subpopulations that differ in phenotype, surface marker expression, functional capacity, tissue localization, and distribution (7). Conventional DCs (cDCs) are potent antigen presenting cells that prime T cells and initiate adaptive response; plasmacytoid DCs (pDCs) are characterized by a fast and abundant type I interferon response, particularly in response to viral or nucleic acid stimuli (12). Monocyte-derived DCs (moDCs) arise from circulating monocytes in inflammatory conditions and promote T-helper responses; in contrast tolerogenic DCs (tolDCs) exhibit lower levels of co-stimulatory molecules and higher anti-inflammatory signals, promoting immune tolerance and regulatory T cell activity (11,13,14). Dysregulated DCs subsets, altered localization, and frequencies have been widely associated with SLE pathogenesis (7,10,14,15).

DCs are present in peripheral blood and constitute less than 1% of the circulating peripheral blood mononuclear cells, which makes their direct recovery for research purposes challenging (16). As a result, *in vitro* models such as moDCs have been widely used. Although moDCs are not fully equivalent to *ex vivo* DCs, they display hallmark functional features that make them a widely accessible *in vitro* system for studying DCs properties and functions (7). In particular, moDCs can exhibit strong proinflammatory activity and promote inflammatory T helper responses (Th1, Th2, and Th17), reflecting key aspects of DCs mediated immune activation, which makes them useful for studying mechanisms relevant to inflammation and autoimmune conditions (10,15).

TolDCs have the capacity to inhibit self-reactive responses by modulating T cell activity, inducing T cell anergy, or supporting regulatory T cell (Treg) generation. They are characterized by low expression of co-stimulatory molecules (e.g., CD80, CD86, and CD83), proinflammatory cytokines and major histocompatibility complex (MHC) molecules, and elevated anti-inflammatory markers (e.g. IL-10, TGF-β, PD-L1, and PD-L2) (7,10,11,14). Over the last decades, several efforts have focused on characterizing tolDCs and their capacity to restore tolerance as a therapeutic approach for autoimmune diseases such as type 1 diabetes (T1D) mellitus, rheumatoid arthritis (RA), multiple sclerosis (MS), and Crohn’s disease (17). When treated with dexamethasone and rosiglitazone, moDCs from SLE patients adopted a tolerogenic phenotype with promising functional capacity to suppress T cell activation, with no significant differences found between SLE patients and healthy controls (18).

Nevertheless, comparing DCs subsets between SLE patients and healthy controls at a single differentiation endpoint is unable to capture the nuanced alterations in monocytes to DCs differentiation programs. Therefore, a more detailed analysis of how transcriptional trajectories and underlying regulatory networks diverge during the generation of moDCs and tolDCs is required to reveal disease-specific biology. In this study, we performed transcriptomic profiling of moDCs and tolDCs derived from Mexican women with SLE and healthy controls to dissect these differentiation processes, identify alterations in tolerance related pathways, and highlight potential mechanistic targets in SLE pathophysiology.

## Methods

### Ethics

This study was conducted in accordance with the Declaration of Helsinki and approved by the Ethics on Research Committee of the Instituto de Neurobiología at the Universidad Nacional Autónoma de México (UNAM, Protocol 093.H), and Instituto Nacional de Ciencias Médicas y Nutrición Salvador Zubirán (IRE-3636). Written informed consent was obtained from all participants before enrollment. All clinical and demographic data was anonymized and securely stored at the Laboratorio Nacional de Visualización Científica Avanzada (LAVIS), UNAM, to ensure its confidentiality.

### Participant recruitment and sample collection

Mexican women aged between 18-50 years with SLE were identified through the Mexican Lupus Registry (LupusRGMX), part of the MEXOMICS consortium, and by certified rheumatologists (19,20). SLE diagnosis was confirmed using the 2019 European League Against Rheumatism/American College of Rheumatology Classification Criteria (EULAR/ACR). In this analysis, only participants with glucocorticoid therapy at <10 mg/day were included; however, information on the duration of the use of the current dose was not collected. We confirmed that there had been no changes in treatment over the past two months and that no biological treatment had been used in the previous six months. Exclusion criteria included pregnancy, additional chronic diseases (including other autoimmune diseases), concurrent hormonal or antibiotic treatments, use of biological treatments, and/or any modification of SLE-specific therapy in the previous two months.

In general, SLE participants were evaluated by rheumatologists, a subset of cases relied on participant self-report supported by review of their personal medical records (e.g., laboratory results and clinical notes), as electronic medical records are not consistently available in Mexico.

EDTA-anticoagulated whole blood was obtained by peripheral venous puncture from 23 Mexican women with SLE and 10 healthy controls from June 2022 to February 2023. The clinical and sociodemographic characteristics of all volunteers are summarized in Supplementary Table S1.

### Generation of moDCs and tolDCs

Peripheral blood mononuclear cells (PBMCs) were recovered using Lymphoprep™ (Stemcell Technologies) density gradient. Briefly, blood was diluted 1:1 (v/v) in PBS 1X, gently poured over Lymphoprep™ (Stemcell Technologies) and centrifuged at 3000 rpm, 20 minutes, at room temperature, without break. PBMCs were collected from the interface formed at the density gradient. Monocytes were then isolated from PBMCs using the negative selection EasySep™ Human Monocyte Isolation Kit (Stemcell Technologies) magnetic approach according to the manufacturer’s instructions. Cells were then recovered for RNA extraction and flow cytometry.

Purified monocytes were resuspended in RPMI-1640 medium supplemented with 10% FBS and seeded at a density of 1.5×10^6^ cells per 2mL in 24-well plates. For each donor, two wells were prepared: one for monocyte-derived dendritic cells (moDCs) and another for tolerogenic dendritic cells (tolDCs) differentiation. Initially, both wells were supplemented with 1 ng/mL of recombinant human granulocyte-macrophage colony stimulating factor (GM-CSF) (Preprotech) and 25 ng/mL of recombinant human interleukin-4 (IL-4) (Preprotech). For moDCs, cultures followed the supplementation with GM-CSF and IL-4 each 48 hours. For tolDCs, cells undergo the same supplementation plus a daily dose (40 ng/mL) of recombinant human interleukin 10 (IL-10) (Preprotech) from day 2 onward. On day 8, cells were harvested for flow cytometric analysis and RNA extraction.

### Flow cytometry and gating strategy

The expression of surface markers associated with monocyte and DCs antigen presentation mechanisms was measured in 6 SLE and 4 control samples using flow cytometry. Cells were initially incubated with Human Fc Receptor blocking solution (TruStain FCX™, BioLegend) for 10 minutes, and then incubated with a cocktail of fluorochrome-conjugated antibodies: anti-human CD14 [myeloid cell surface marker] redFluor 710 (Clone 61D3, Tonbo Biosciences), anti-human CD11c [dendritic cell surface marker] FITc (Clone 3.9, Invitrogen), anti-human HLA-DR [dendritic cell surface marker] APC (Clone L243, Tonbo Biosciences), anti-human CD40 [dendritics cell co-stimulatory molecule] PE (Clone G28.5, Tonbo Biosciences) and anti-human CD80 [dendritics cell co-stimulatory molecule] PerCP-Cy 5.5 (Clone 2D10, Tonbo Biosciences). Staining was performed for 30 minutes at 4°C, in the dark.

The gating strategy is detailed in Supplementary Figure S1. Briefly, we gated on the total cell population by forward scatter area (FSC-A) versus side scatter area (SSC-A) to exclude debris. Singlets were selected by gating on FSC-H vs. FSC-A to remove doublets, as recommended in flow cytometry guidelines. Within the singlet population, cells positive for both CD14 and CD11c were identified, defining the myeloid/monocyte-DCs compartment. We then measured the expression of HLA-DR, CD40, and CD80 within this CD14^+^CD11c^+^ gate to assess DCs maturation and antigen-presenting potential. Data was recovered and analyzed using FlowJo v7.6.2, and visualized in R using the ggplot2 package.

### RNA isolation and library preparation

Recovered cells were resuspended in 200 μL of RNAlater and stored at −20°C up to extraction. Cells were resuspended in 1 mL of PBS 1X and centrifuged for 7 minutes at 14,000 rpm and 4°C, and then resuspended in 500μL of Trizol reagent (TRIzol® RNA Isolation Reagent, Invitrogen) and stored at −70°C for the night. Total RNA was extracted using RNeasy Mini Kit (Qiagen) according to the manufacturer’s instructions. Samples were treated with DNase I (TURBO DNA-free™ Kit Invitrogen) to eliminate any residual DNA. RNA concentration, quality and integrity was assessed using the RNA high sensitivity kit (Qubit™ RNA HS Assay Kit, Invitrogen) and the Qubit 4 Fluorometer (Invitrogen). Samples were stored in RNAse-free water at −20°C until sequencing.

RNA sequencing was performed by Novogene Inc. (Sacramento, CA, United States) using their mRNA-seq service. Libraries were prepared by poly A enrichment, and sequencing was performed on the Illumina NovaSeq X Plus® system in a 150 bp paired-end format. Details on all transcriptomes generated in this study are provided in Supplementary Table S2.

### Differential expression analysis

Bulk RNA-seq data was analyzed using Nextflow (v23.04.3) in combination with Singularity (v3.7.0). Preprocessing of raw data was performed with the nf-core/RNAseq pipeline (v.3.14.0; https://github.com/nf-core/rnaseq, SciLifeLab) using GRCh38/hg38 (GCA_000001405.2) downloaded from UCSC as reference genome. Quality control and adapter trimming were carried out with Trim Galore! (https://github.com/FelixKrueger/TrimGalore), while alignment and transcript quantification were conducted with the STAR–Salmon workflow (21).

Quantification results from Salmon were imported into R using the tximport package (v1.18.0) (22). To explore global variations in gene expression profiles between the experimental groups (SLE and control), raw counts were transformed using a variance stabilization approach (VST) implemented in the DESeq2 package (v1.30.1) (23) and Z-score normalized. A Principal Component Analysis (PCA) was applied to the normalized expression matrix, and the first two principal components were selected for visualization (Supplementary Figure S3).

To identify differentially expressed genes (DEGs), the DESeq2 package (v.1.30.1) (23) was used on the raw count matrix, applying the Wald test for pairwise comparisons. Multiple testing correction was conducted by controlling the false discovery rate (FDR) at 0.05 using the Benjamini–Hochberg (BH) procedure (24). We defined differentially expressed genes as those with an absolute log2FoldChange (LFC) greater than 0.5 and an adjusted p-value (padj) < 0.05. To address our different questions, the following pairwise comparisons were applied:

1. **Comparisons between SLE vs. controls within each cell type:** DEGs were identified by comparing SLE patients to healthy controls separately in each cell type (monocytes, moDCs, and tolDCs). This approach allowed the identification of cell-type-specific transcriptional signatures associated with SLE. The model employed was ∼ Group in each cell type.
2. **Comparison of monocyte-to-DC differentiation trajectories between SLE and non-SLE controls:** We characterized transcriptional changes associated with monocyte-to-dendritic cell differentiation, independent of disease group. moDCs and tolDCs were compared to monocytes separately within the SLE and control groups. This analysis provided baseline differentiation signatures in healthy controls and revealed how these programs were altered in SLE patients. The model employed was ∼ Group + Cell_type.
3. **SLE interaction effect on the moDCs and tolDCs differentiation:** This interaction analysis highlighted SLE-specific effects on monocyte-to-DCs transcriptional programs. The model employed was ∼ Group + Cell_type + Group:Cell_type.

To identify differentially expressed genes shared with the differentiation analysis results (comparison 2) or unique to this interaction dataset, we merged the tables using the full_join or left_join function from the dplyr package in R (v1.1.4).

### Gene Ontology and Pathway Enrichment Analysis

For each set of differentially expressed genes of interest, functional enrichment analysis was performed using g:Profiler2 (v0.2.0) (25), together with supporting R packages: enrichplot (v1.10.2) (26), DOSE (v3.16.0) (27), clusterProfiler (v3.18.1) (28,29), and tidyverse (v2.0.0) (30). Genes were queried against multiple annotation databases, including Gene Ontology (GO: Biological Process, Cellular Component, Molecular Function), Reactome (REAC), Human Phenotype Ontology (HP), KEGG, WikiPathways (WP), transcription factor (TF) targets, CORUM protein complexes, and miRNA annotations. Significance was assessed using the hypergeometric test, with multiple testing corrections applied through the FDR and BH procedure. Only terms with padj < 0.05 were considered significant.

### Regulatory network analysis

To explore gene regulatory mechanisms, we used pySCENIC’s (https://github.com/aertslab/pySCENIC) “scenic_multiruns” pipeline implemented by VSN-Pipelines (v0.25.0) (31), executed with Nextflow (v23.04.3) in combination with Singularity (v3.7.0). This enabled the construction of protein–protein interaction, co-expression, and competitive endogenous RNA (ceRNA) networks, as well as to infer regulatory modules. The input was the VST-transformed gene expression count matrix obtained using DESeq2 (v.1.30.1) (23). We filtered for regulons (sets of transcription factors and their targets) that were found in at least two runs across all iterations, and target genes had to be assigned to the same transcription factors (TF) at least five times.

Regulon activity matrices of AUC (area under the recovery curve) values were extracted from the resulting SCENIC’s loom file and matched to the corresponding sample metadata. To determine differential regulons, we first evaluated overall AUC normality using the Shapiro–Wilk test. Regulons with normally distributed AUC values were tested using an unpaired Student’s t-test, whereas non-normal regulons were evaluated with a Mann–Whitney U test. Multiple testing correction was done using the Benjamini–Hochberg method in Python (v3.8.19) and the SciPy package (v1.10.1). Log₂fold change (log_2_FC) was computed per regulon from sample-wise mean AUC values. We deemed a regulon differential if the adjusted p-value was < 0.05 and the absolute log_2_FC was > 0. The analyses were performed following the steps outlined below:

1. To identify regulons significantly altered in SLE, we compared AUC values from SLE samples to those from control samples within each cell type, using controls as the reference group.
2. To assess changes along the monocyte-to-DC differentiation trajectories, we compared moDCs AUC values to those of monocytes, and tolDCs AUC values to those of monocytes, using monocytes as the reference in both SLE and control samples. Then we subset the target genes of SLE-exclusive regulons that also showed differential expression in comparison 2.

To identify patterns that could be shaping the global behavior of SLE-derived moDCs and tolDCs separate from controls, we first subset the AUC values (z-score normalized) to the regulons that target shared DEGs between moDCs and tolDCs, from the interaction analysis (comparison 3). For each regulon *j*, we removed the effect of the cell type (moDCs or tolDCs) by fitting a linear model of the form:

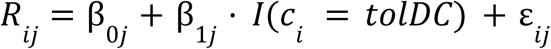

Where:

- *R_ij_* denotes the regulon *j* activity (Z-score) for sample *i*,
- *B_0j_* is the mean regulon activity on moDCs,
- *B_1j_* represents the average difference between tolDCs and moDCs,
- *I*(·) is a binary indicator variable equal to 1 for tolDCs and 0 for moDCs.
- *c_i_* represents the cell type *c* for each sample *i*.
- and *ε_ij_* represents the residuals.

To reveal variation not explained by cell type, we extracted a matrix of residuals and performed PCA on those values. Then, to evaluate each regulon’s contribution, we compared the absolute mean Z-scores between SLE and controls, prioritizing the regulon with the largest absolute mean difference.

### Statistical analyses and data visualization

Data and statistical analyses of the clinical and sociodemographic data were performed using R (v.4.0.2) (32). The normal distribution of sociodemographic and clinical data was evaluated using the Shapiro-Wilk test. For variables with normal distribution, the mean and standard deviation (SD) were reported, while for variables with a non-normal distribution, the median, interquartile range (IQR), minimum, and maximum values were calculated. Since age presented a normal distribution, the Student’s t-test was used to compare values between SLE and controls. These tests were performed using the “stats” package (v.3.6.2).

Additional enrichment analyses were evaluated using the hypergeometric over-representation test implemented in phyper, base package in R, and a one-tailed Fisher’s exact test implemented in fisher.test from stats package in R.

All figures including heatmaps, scatterplots, box plots, barplots, networks and upset plots were created using a combination of ggplot2 (v3.5.1) (33), ggh4x (v0.2.8) (34), gridExtra (v2.3) (35), circlize (v0.4.16) (36), reshape2 (v1.4.4) (37), tidyverse (v2.0.0), ComplexHeatmap (v2.20.0) (38) and Cytoscape (39).

## Results

### Overview of the Experimental Workflow

Peripheral blood samples were obtained from 23 Mexican women with SLE and 10 healthy controls recruited through LupusRGMX and certified rheumatologists. All SLE diagnoses were confirmed using the 2019 EULAR/ACR classification criteria, and participants were selected to minimize treatment-related confounding, including stable low-dose glucocorticoid use and absence of recent biologic therapy (See Methods). PBMCs were isolated from each sample, and monocytes were purified for immediate analysis or differentiated *in vitro* into moDCs or tolDCs using established cytokine stimulation protocols (See Methods). Monocytes, moDCs, and tolDCs were then harvested for flow cytometric characterization and bulk RNA sequencing, followed by differential expression, functional enrichment, and regulatory network analyses (**Figure 1**).

**Figure 1.**
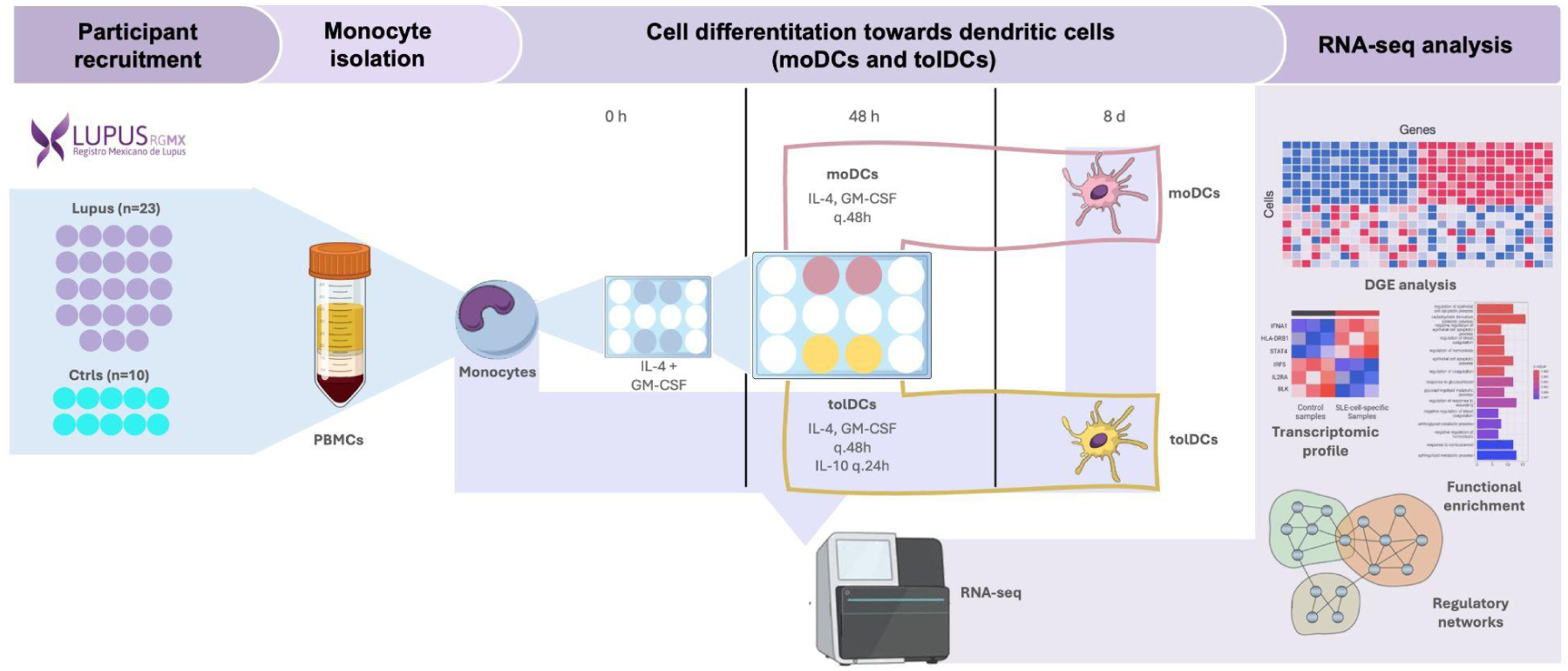
Experimental design of the study. Schematic overview of the experimental workflow. Blood samples were collected from volunteers recruited through LupusRGMX. Peripheral blood mononuclear cells were isolated, and monocytes were subsequently purified and differentiated *in vitro* into moDCs (yellow cells) or tolDCs (pink cells). Monocytes, moDCs and tolDCs were harvested for total RNA extraction, and high-throughput RNA sequencing was performed. Downstream bioinformatic analyses included differential gene expression (DGE), functional enrichment, and regulatory network.

### Clinical Characterization of the Cohort

We initially calculated descriptive statistics and assessed data distribution for all continuous variables (histograms, Q-Q plots, and Shapiro–Wilk tests; see Supplementary Figure S2; Supplementary Table S3), before summarizing sociodemographic and clinical characteristics. We recruited 23 volunteers with SLE and 10 healthy controls from LupusRGMX (sociodemographic and clinical characteristics of the volunteers are presented in Table 1). Volunteers with SLE had a mean age (± SD) of 33.7 (± 8.6) years, whereas the healthy control group had a mean age (± SD) of 29.9 (± 7.0) years. SLE participants reported a median (IQR) age at diagnosis of 27.0 (IQR: 16.0) years, with a median (IQR) of 5.7 (IQR: 4.7) years since the diagnosis. Six (26.1%) of the participants reported a family history of SLE and three (13%) had been diagnosed with lupus nephritis (Supplementary Table S4). Regarding SLE activity, we observed that ten volunteers (43.5%) had disease activity. As for treatment consumption, sixteen (69.6%) of the participants use corticoids on a regular basis, with a median daily dose of 5 mg. The use of antimalarials was also frequently reported by our cohort (78.3%).

**Table 1.** Sociodemographic and clinical characteristics of the volunteers with SLE (n=23) and Ctrl (n=10). SD: Standard Deviation, IQR: Interquartile Range.

| Characteristic | SLE<br>(n=23) | Ctrl<br>(n=10) |
| --- | --- | --- |
| **Age (years)** | Mean±SD: 33.7±8.6 | Mean±SD: 29.9±7.0 |
| **Time with diagnosis (years)** | Median (IQR): 4 (4.5)<br>Min-Max: 1-17 | NA |
| **Age at diagnosis (years)** | Median (IQR): 27 (16)<br>Min-Max: 16-43 | NA |
| **Disease activity (n,%)** |  |  |
| Without activity/low activity (0-5 pts) | 13 (56.5%) | NA |
| With activity (≥6 pts) | 10 (43.5%) |  |
| **Nephritis (n,%)** | 3 (13.0%) | NA |
| **Family history of SLE (n,%)** | 6 (26.1%) | NA |
| **Treatment use (n,%)** |  |  |
| Glucocorticoids use | 16 (69.6%) | NA |
| *Dose (mg/day)* | Median (IQR): 5 (0) |  |
|  | Min-Max: 2.5-8 |  |
| Antimalarials | 18 (78.3%) |  |
| Mycophenolate | 7 (30.4%) |  |
| Azathioprine | 6 (26.1%) |  |
| Metotrexate | 2 (8.7%) |  |
| Rituximab | 1 (4.3%) |  |
| Others | 5 (21.7%) |  |

### Immune Phenotype Variability in SLE-Derived DCs

To evaluate the effects of our different supplementation conditions on cell phenotypes, we performed flow cytometry on all cell types (Supplementary Figure S1). We first quantified the percentage of CD14⁺ [myeloid cell surface marker] and CD11c⁺ [dendritics cell surface marker] cells and found that monocytes had a lower frequency of this population compared to moDCs and tolDCs. Notably, the proportion of CD14⁺CD11c⁺ cells was reduced in the SLE group relative to controls. Next, we measured CD40 [dendritics cell co-stimulatory molecule] expression: all three cell types (mo, moDCs, tolDCs) exhibited similarly high levels, with no differences between SLE and control samples. HLA-DR [dendritics cell surface marker] expression was also high across all samples, however, SLE-derived moDCs showed a slightly broader distribution, which might suggest more heterogeneity of HLA-DR expression in these cells. CD80 [dendritics cell co-stimulatory molecule] also exhibited a high expression among all samples, with SLE samples displaying a slightly wider spread than controls, hinting at greater variability in costimulatory capacity in this group.

To validate the expected cell types, we selected established specific markers previously reported in the literature (Table 2) and assessed the expression of these genes across all samples. We observed the expected upregulation of cell-type–specific markers in their respective cell type samples, confirming transcriptionally that the *in vitro* differentiation protocols were successful (Figure 2B).

**Figure 2.**
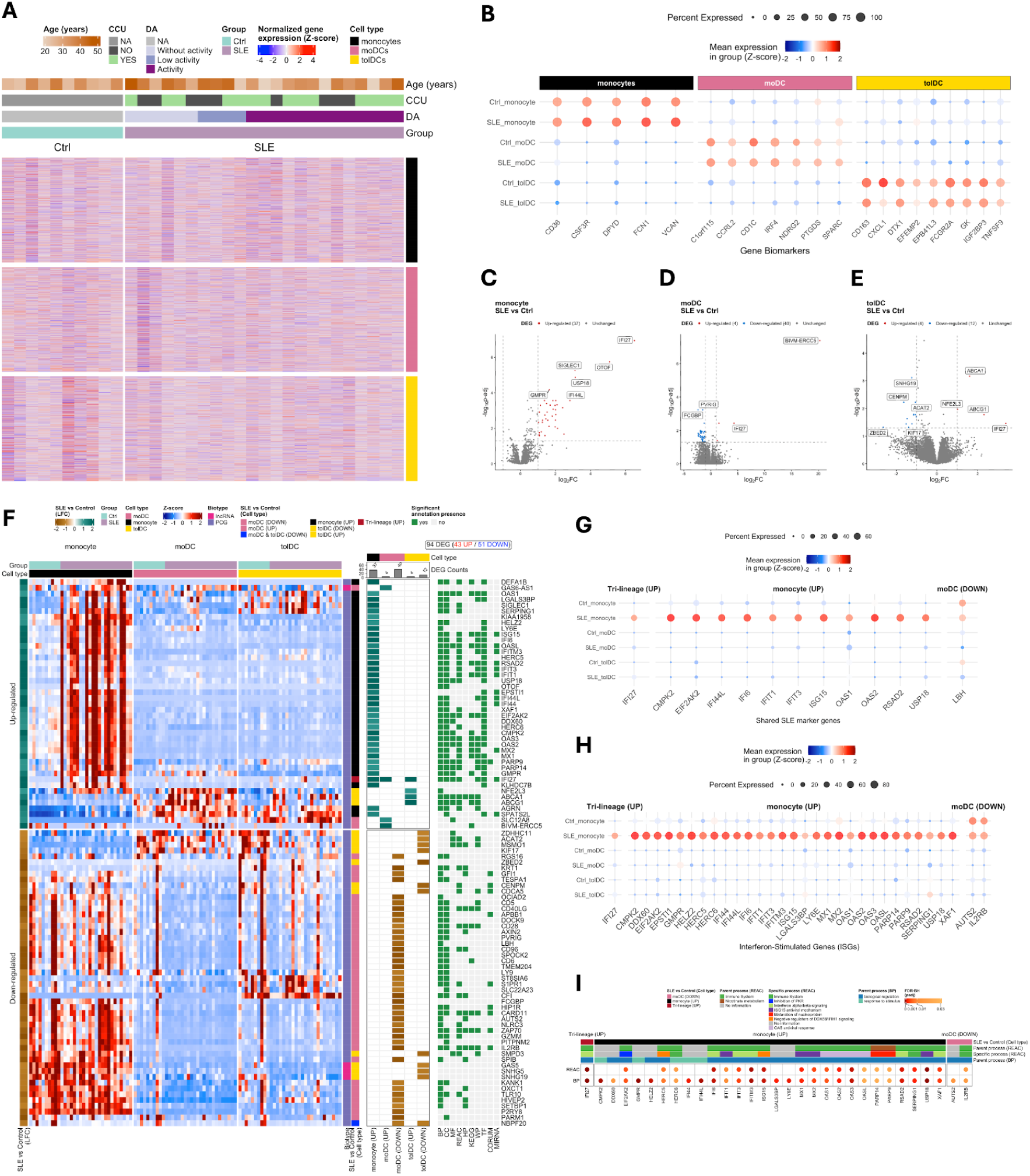
Integrated cellular and transcriptomic profiling across cell types and disease groups reveals cell type specific perturbations and a shared SLE marker. **(A)** Heatmap of normalized gene expression, with genes represented in rows and samples in columns. Samples include mo, moDCs and tolDCs. Color scale represents Z-score-normalized counts. Metadata for each sample is annotated at the top: “Group” denotes the disease status of each sample (SLE patient and Ctrl), “DA” indicates disease activity (or NA for controls), and “CCU” marks corticoid usage. A second annotation highlights the cell type for each sample. **(B)** Cell-type-specific biomarkers. Color scale indicates average expression per group (z-score normalization); dot size reflects the percentage of samples expressing each marker. Each column is a different biomarker gene for the cell types annotated at the top. Volcano plots of differentially expressed genes in **(C)** mo, **(D)** moDCs, and **(E)** tolDCs between groups, selection was based on LFC ≥ 0.5 and padj < 0.05, using the Wald test with Benjamini-Hochberg correction for FDR. **(F)** Integrated visualization of DEGs and functional enrichment in SLE. **Left panel:** Heatmap of 57 DEGs (26 upregulated and 31 downregulated) between SLE and Ctrl groups across cell types, selected using LFC ≥ 0.5 and padj < 0.05. Color scale corresponds to Z-scores. **Middle panel:** Heatmap showing log₂FoldChange values for SLE versus Ctrl (brown to green) across cell types. Each column includes a barplot indicating the number of DEGs per cell type: 20 mo (up), 4 moDCs (up), 29 moDCs (down), 4 tolDCs (up), and 12 tolDCs (down). **Right panel:** Heatmap of functional terms enriched among DEGs, identified using g:Profiler2 across multiple databases (GO:BP, CC, MF, REAC, HP, KEGG, WP, TF, CORUM, and miRNA). Color scale indicates presence or absence of significant annotation based on hypergeometric testing. Panels **(G)** and **(H)** share the same color scale, indicating average expression (Z-score normalization) per disease group-cell type (rows). Gene markers are in the columns, classified in three groups according to their shared expression patterns across cell types: “Tri-lineage” denotes genes upregulated in monocytes, moDCs, and tolDCs. “Monocyte (UP)” refers to genes consistently upregulated in monocytes, while “moDCs (DOWN)” indicates genes consistently downregulated in moDCs. **(G)** SLE marker genes and **(H)** Differentially expressed ISGs (DEISG); dot size reflects the percentage of samples expressing each marker. **(I)** Heatmap of biological processes associated with the DEGs. Processes were classified according to Reactome (REAC) parent and specific categories, and Gene Ontology Biological Process (GO:BP) parent terms. REAC parent processes include Immune system, Synthesis of phosphatidylinositol (PI), Cell division, Lipid metabolism and Transport of small molecules. Color scale represents padj (FDR–BH correction), with a maximum threshold of padj < 0.05. SLE: Systemic Lupus Erythematosus; Ctrl: healthy control; mo: monocytes; moDCs: monocyte-derived dendritic cells; and tolDCs: tolerogenic dendritic cells; DEGs: differential gene expression; padj: adjusted p-value; LFC: log₂FoldChange; FDR: false discovery rate; BH: Benjamini–Hochberg procedure.

**Table 2.** Genetic markers of cell identity.

| Cell type | Genes associated | References |
| --- | --- | --- |
| Monocytes | *CD36* | (40) |
|  | *CSF3R* | (41) |
|  | *DPYD* | (42) |
|  | *FCN1* | (43) |
|  | *VCAN* | (42–44) |
| moDCs | *C1orf115* | (45) |
|  | *CCRL2* | (46) |
|  | *CD1C* | (47–49) |
|  | *IRF4* | (48,49) |
|  | *NDRG2* | (43) |
|  | *PTGDS* | (50) |
|  | *SPARC* | (51) |
| tolDCs | *CD163* | (45) |
|  | *CXCL1* | (14) |
|  | *DTX1* | (52) |
|  | *EFEMP2* | (14) |
|  | *EPB41L3/FBLN4* | (45) |
|  | *FCGR2A/CD32* | (43) |
|  | *GK* | (45) |
|  | *IGF2BP3* | (45) |
|  | *TNFSF9/4-1BBL/CD137L* | (53,54) |

### Cell-Type–Specific Transcriptional Signatures of SLE

To understand the general differential expression signature of SLE in monocytes, moDCs and tolDCs, we generated transcriptomic data for each cell type (Figure 2A). PCA of normalized expression confirmed the separation between cell types and the absence of batch effects (variance explained by PC1: 62.23%, PC2: 9.33%) (Supplementary Figure S3). We performed differential expression analysis between SLE and controls for each cell type. In total, we identified 94 differentially expressed genes (DEGs) across all comparisons (Supplementary Table S5; Supplementary Table S6). These comprised: 37 DEGs in monocytes, all of them upregulated in SLE (Figure 2C); 44 in moDCs (4 upregulated and 40 downregulated) (Figure 2D); and 16 in tolDCs (4 upregulated and 12 downregulated) (Figure 2E). Their expression patterns are shown in Figure 2F, highlighting cell-type-specific expressions.

Interferon-stimulated genes (ISGs) are central drivers of monocyte and dendritic cell activation, and their persistent upregulation is a defining feature of SLE (54). Hence, we set out to identify ISGs within our DEG using a compiled list (55) (Figure 2H; Supplementary Table S8). ISGs were overrepresented among DEGs (hypergeometric test, p-value = 2.48 × 10^-31^; Fisher’s exact test, p-value < 2.2 × 10^-16^). Among all ISGs, *IFI27* was the only gene up-regulated across all three cell types in SLE (denoted as Tri-lineage in Figure 2G; Figure 2H), underscoring its potential relevance in SLE pathophysiology (56–58).

Specifically, monocytes from SLE patients exhibited 30 differentially expressed ISGs (DEISGs)(Figure 2H; Supplementary Table S8). Among these, differences in *IFI27*, *IFI6*, *IFI44L* and *USP18* expression are consistent with previous reports, supporting their strong potential as biomarkers to distinguish SLE from controls; in addition, the marked enrichment of these ISGs in monocytes highlights a potentially central role of monocytes in sustaining lupus-associated transcriptional programs (57–63). In contrast, among moDCs we found two ISGs that had differential gene expression between SLE and healthy controls (AUTS2 and *IL2RB/CD122*), and no ISGs were significant for tolDCs (Figure 2H). Replicating results previously showing no significant differences found between SLE patients and healthy controls (18).

Established SLE transcriptional signatures have been widely reported in prior studies. Hence, we integrated a reference list of SLE-associated genes (Supplementary Table S7) used to assess the disease relevance of our findings. We observed a significant enrichment of SLE-associated genes among the set of DEGs (hypergeometric test, p-value = 1.144 × 10^-11^; Fisher’s exact test, p-value = 1.144 × 10^-11^) (Figure 2G; Supplementary Table S7). Specifically, monocytes from SLE patients showed elevated expression of *IFI27*,*CMPK2, EIF2AK2, IFI44L, IFI6, IFIT1, IFIT3, ISG15, OAS1, OAS2, RSAD2* and *USP18,* all of them well-established SLE-associated genes (Figure 2G). In contrast, among moDCs we found downregulation of one gene previously associated with SLE: *LBH* (Figure 2G); and no SLE-associated genes were significant for tolDCs. These findings point to cell-type–specific transcriptional shifts that may alter DCs signaling and immune regulation in SLE.

Pathway enrichment analysis of DEGs revealed immune-related Reactome (REAC) pathways (Figure 2I; Supplementary Table S8). Among monocyte DEGs, the most frequent parent process (BP) was immune system related processes, among which specific process (REAC) included: interferon alpha/beta signaling emerged as the most significantly enriched pathway (*IFI5, IFIT3, IFITM3, RSAD2, XAF1*), followed by ISG15 antiviral mechanism (*IFIT1, MX1, MX2, USP18*), and OAS antiviral response (*OAS1, OAS2, OAS3, OASL*) and maturation of nucleoprotein (*PARP14* and *PARP9*) (Supplementary Figure S4). Interferon signaling plays a decisive role in reshaping monocyte transcriptional programs in SLE, a process that likely underpins the systemic inflammatory and autoimmune manifestations characteristic of the disease (55,56,62). For moDCs, *IL2RB* was associated with the immune system, whereas *AUTS2* was not found to be associated with any specific process. Regarding tolDCs, no ISGs or SLE-associated genes were observed.

In order to investigate the regulatory layer behind these DEGs, we assessed whether regulon activity differed between SLE and control samples within each DCs type, using AUC-based regulon scores and the statistical framework described above (See Methods). After multiple-testing correction (LFC > 0 and padj < 0.05), no regulons reached statistical significance in any of the cell types analyzed. This suggests that, at the bulk level for these specific cell types, transcriptional regulatory changes between SLE and controls are subtle and do not manifest as robust, global shifts in regulon activity, confirming previous findings (18).

### Immune and metabolic remodeling during moDCs and tolDCs differentiation in SLE

Our initial analyses revealed that most transcriptional variation was attributable to cell-type identity, with minimal differences between SLE and controls; this was also reflected in the regulon profiles (Supplementary Figure S5) in which the major source of variation corresponds to cell type instead of condition (Supplementary Figure S6), and tolDCs showed no disease-specific signals. Consequently, we focused on comparing the differentiation trajectories themselves, assessing how the monocyte-to-moDCs and monocyte-to-tolDCs transitions differed between SLE and healthy individuals (Figure 3A). In total, we found over 8,114 unique DGEs whose expression levels changed significantly across these comparisons (4427 upregulated and 3752 downregulated). We found: 2,688 up- and 2,947 downregulated DEGs in monocytes vs moDCs from controls; 2,736 up- and 2,851 downregulated DEGs in monocytes vs tolDCs from controls; 2,945 up- and 3,652 downregulated DEGs in monocytes vs moDCs from SLE; and 2,804 up- and 3,591 downregulated DEGs in monocytes vs tolDCs from SLE (Figure 3B; Supplementary Table S9).

**Figure 3.**
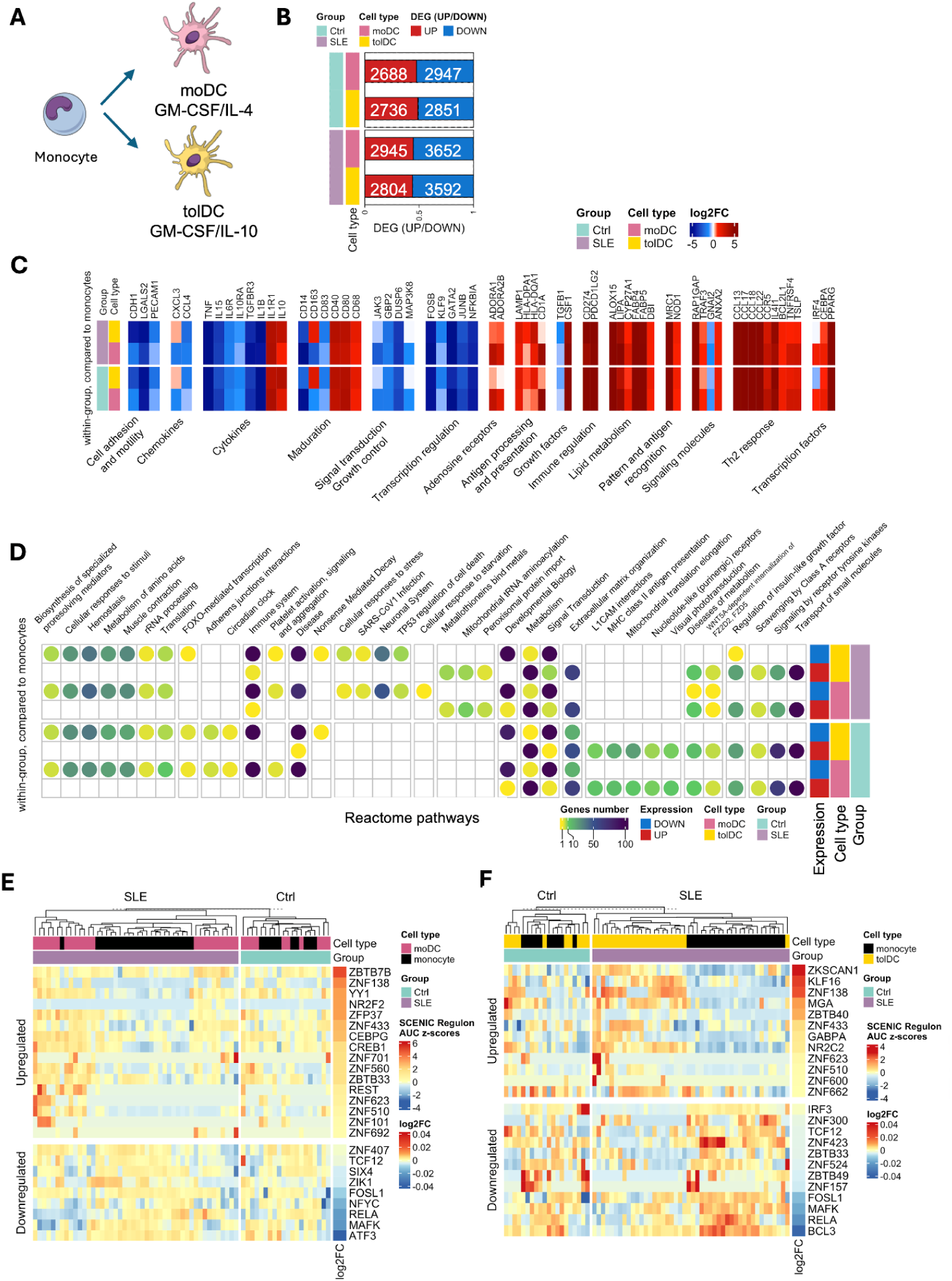
Monocyte differentiation to moDCs and tolDCs. **(A)** Schematic representation of monocyte differentiation into DCs (moDCs and tolDCs). **(B)** Number of differentially expressed genes (DEGs) in DCs of both SLE and controls, using monocytes as reference. **(C)** Genes previously reported as up- or downregulated during moDCs differentiation compared with monocytes, showing the expected transcriptional changes in our dataset. D) Biological processes enriched among DEGs. **(E)** Exclusive regulons associated with monocyte differentiation into moDCs. **(F)** Exclusive regulons associated with monocyte differentiation into tolDCs.Significance thresholds were set at absolute LFC ≥ 0.5 and padj < 0.05. SLE: Systemic Lupus Erythematosus; Ctrl: healthy control; mo: monocytes; moDCs: monocyte-derived dendritic cells; and tolDCs: tolerogenic dendritic cells; DEGs: differential gene expression; padj: adjusted p-value; LFC: log₂FoldChange; FDR: false discovery rate; BH: Benjamini–Hochberg procedure.

Gene expression studies of monocyte differentiation into moDC s have consistently identified characteristic transcriptional changes that define a differentiation signature. A hallmark of this process is the downregulation of classical monocyte markers, including CD14 and CD163 (14,42). In our analysis, we confirmed that CD14 was repressed in both moDCs and tolDCs, in agreement with earlier studies in moDCs (14). However, CD163 displayed a distinct pattern: while downregulated in moDCs as expected, it was upregulated in both tolDCs from SLE and controls (Figure 3C; Supplementary Table S10). In addition, we observed overexpression in tolDCs differentiations (both SLE and controls) of genes related to DCs maturation including *CD40, CD80* and *CD68*, and with the production of anti-inflammatory cytokines, such as *IL-10* and *IL1R1* (Figure 3C; Supplementary Table S10) which are required for tolerance induction (57). Furthermore, moDCs and tolDCs from both SLE patients and controls showed significant downregulation of *CD83*, a marker associated with DCs maturation (58,60,63) (Figure 3C). Consistent with established differentiation signatures in DCs.

Consistent with previous reports, we found that moDCs exhibited upregulation of genes associated with recognition and antigen uptake (*MRC1* and *NOD1*), antigen processing and presentation (*LAMP1, HLA-DPA1, HLA-DQA2* and *CD1a*), growth factors (*CSF1*), cytokines and their receptors (*IL-10* and *IL1R1*), Th2 response (*CCL13/MCP-4, CCL17/TARC, CCL18/PARC, CCL22/MDC, CCR5, IL4I1, BCL2L1, TNFRSF4* and *TSLP*), lipid metabolism (*ALOX15, LIPA, CYP27A1, FABP4* and *FABP5*), adenosine receptors (*ADORA1* and *ADORA2B*), signaling molecules (*RAP1GAP, TRAF3,* and *ANXA2*), and transcription factors (*IRF4, CEBPA/C/EBP–α,* and *PPARG/PPAR-γ*) (Figure 3C). In contrast, we observed downregulation of genes associated with cell adhesion and motility (E-cadherin/CDH1, galectin-2/LGALS2 and PECAM1/CD31), chemokines (CXCL3/MIP-2β and CCL4/MIP-1β), cytokines and cytokine receptors tumor necrosis factor (TNF)-α (TNF, IL-15, IL-6R, IL10RA and TGFBR3), signal transduction/growth control (JAK3, GBP2, DUSP6, and MAP3K8), and transcriptional regulators (FOSB, KLF9, GATA2, JUNB and NFKBIA) (Figure 3C). Notably, in contrast to earlier studies, we also observed downregulation of *TGFB1, GNAI2,* and *IL-1β*, which had previously been reported as upregulated in moDCs (14,61).

A similar expression pattern was observed between differentiation trajectories in moDCs and tolDCs, with two major exceptions: *MAP3K8* was not differentially expressed in tolDCs comparisons, whereas *CXCL3* was consistently upregulated in tolDCs, while downregulated in moDCs (Figure 3C). Specifically, in tolDCS, the overall expression pattern was maintained, with the exception of IRF4, which was downregulated only in controls showing no significant difference in SLE (Figure 3C).

Pathway enrichment analysis of DEGs revealed distinct patterns associated with monocyte to moDCs differentiation for both SLE and controls (Figure 3C). In controls, we observed downregulation of pathways including: FOXO-mediated transcription, adherens junctions interactions, extracellular matrix organization and circadian clock regulation (Figure 3D). Conversely, upregulation of pathways associated with L1CAM interactions, MHC class II antigen presentation, mitochondrial translation elongation, nucleotide-like purinergic receptors and visual phototransduction was observed. For only SLE, several pathways were downregulated, including: cellular response to stress, SARS-CoV-1 infection, neuronal system, TP53 regulates transcription of cell death gene and cellular response to starvation. In contrast, we found upregulation of metallothioneins bind metals, mitochondrial tRNA aminoacylation, and peroxisomal protein import.

Pathways involved in the monocyte to tolDCs differentiation for controls included the upregulation of L1CAM interactions, MHC class II antigen presentations, mitochondrial translation elongation, nucleotide-like purinergic receptors, visual phototransduction, and disease. In contrast, downregulated pathways among tolDCs controls included: extracellular matrix organization. SLE tolDCS showed upregulation of pathways associated to metallothioneins binds metals, mitochondrial tRNA aminoacylation, and peroxisomal protein import, with downregulation of regulation of insulin-like growth factor (IGF), cellular response to stress, SARS-CoV-1 infection, neuronal system and TP53 regulates transcription of cell death gene (Figure 3D).

To understand the regulatory mechanisms particularly driving the differentiation process to DCs phenotypes, we built gene regulatory networks (see Methods). We identified 32 upregulated and 58 downregulated regulons (absolute log_2_FC > 0 and adjusted p-value < 0.05) upon differentiation to moDCs (Figure 3E), that were common between SLE and controls. We identified 16 upregulated and 9 downregulated SLE-specific regulons during differentiation into moDCs. In contrast, we found only 1 downregulated regulon (*IZNF524*) that was unique to the control samples in this same process, and no control-specific upregulated regulons. In the differentiation towards tolDCs, we identified 26 upregulated and 63 downregulated regulons in common between SLE and healthy controls (Figure 3F). When inspecting SLE-specific regulons, we found 12 upregulated and 12 downregulated. We did not find any control-specific upregulated regulons but we did find 2 downregulated ones: *IRF4* and *YY1*.

To further examine the relevance of these regulons in SLE, we analyzed the target genes of regulon TFs that were uniquely associated with SLE during differentiation into DCs phenotypes. For this, we subsetted those that were also found to be differentially expressed. We found that upregulated and downregulated regulons in the tolDCs differentiation process were composed mostly of downregulated genes (LFC < 1 in DE) (Supplementary Figure S7). We note that among downregulated regulons, ZBTB33 regulates *EIF2AK2*, which has been found as part of the SLE-Related Monocyte signature (SLERRA signature) (55)*. EIF2AK2* was found in our DE analysis as downregulated with LFC = −1.97. In addition, the regulons ZBTB33 and ZBTB7B, which were found upregulated upon moDCs differentiation, also regulate SLERRA signature genes (*EIF2AK2* by ZBTB33 and *LY6E* by ZBTB7B, DE LFC = −2.07 and −1.66, respectively).

We next conducted gene enrichment analysis of the target genes of the significant regulons involved in the differentiation of both dendritic cell phenotypes. We found that in moDCs, neuron development and cell differentiation processes were enriched for genes targets of the upregulated regulons. In contrast, the targets for regulons upregulated in tolDCs were enriched in terms related to transmembrane transport and lymph node development. More interestingly, we found the pathway “DCC mediated attractive signaling” to be associated with gene targets for downregulated regulons in tolDCs, these genes were ABLIM1 (DE LFC = −1.58), ABLIM3 (DE LFC = −1.04) and NCK1 (DE LFC = 1.27), regulated by ZNF157, ZBTB49 and TCF12, respectively (Supplementary figure S8).

Lipid metabolism and associated mechanisms have been reported to be involved in the pathogenesis and progression of SLE (55). Hence, we explored the contribution of previously identified DEGs (64) associated with lipid metabolism to the differentiation of monocytes into moDCs and tolDCs. We detected 56 differentially expressed genes (DEGs) in moDCs and tolDCs from SLE patients compared with controls, which are associated with β-oxidation, fatty acid synthesis, and desaturation pathways, including members of the ACOT family, ACSL3, FASN, ACACA, ACOX2/3, HADH, and HADHB (Figure 4A).

**Figure 4.**
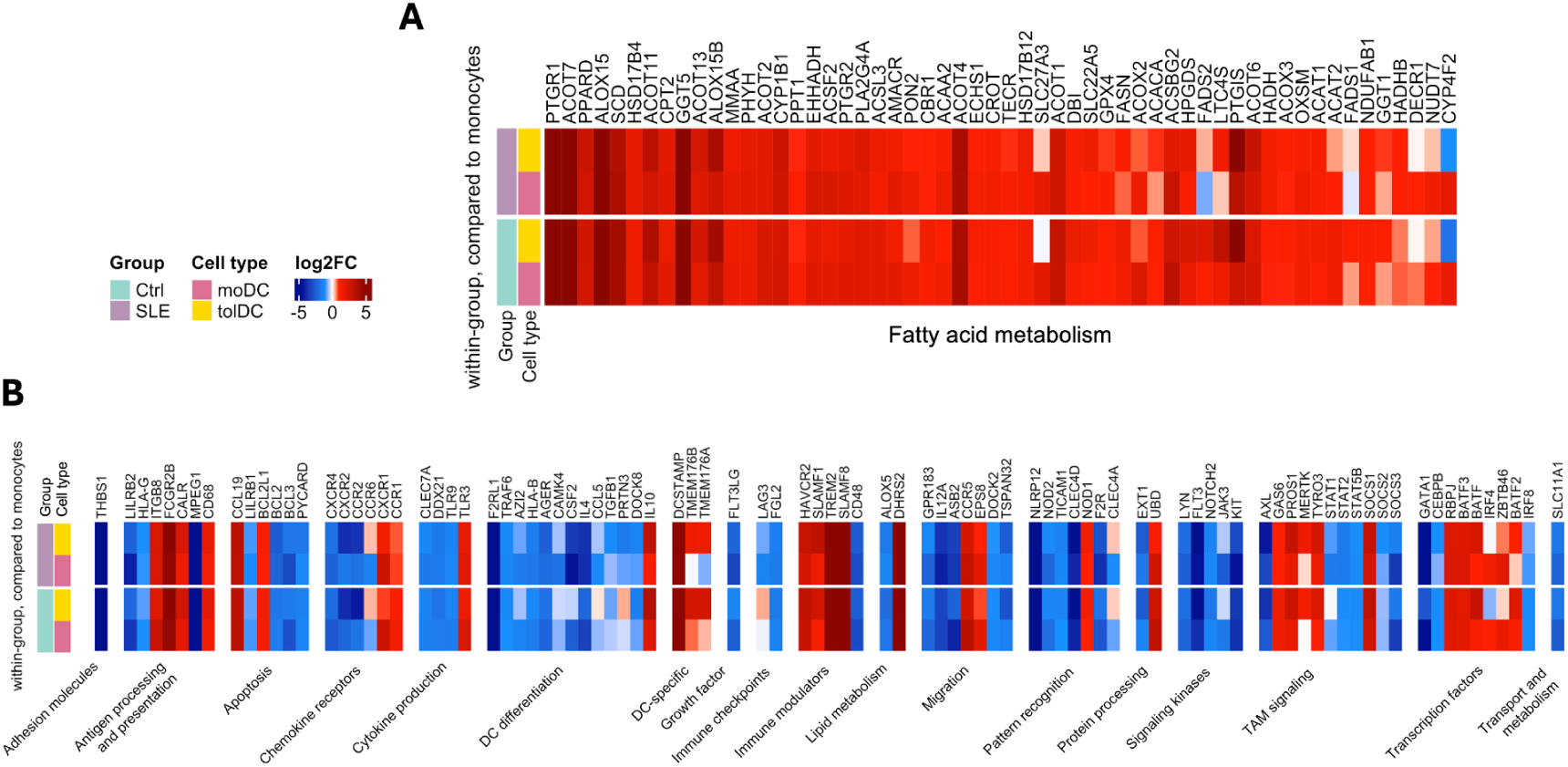
Differentially expressed genes (DEGs) between dendritic cells and monocytes from SLE patients and healthy controls, related to DCs differentiation. (A) Genes associated with fatty acid metabolism, reflecting metabolic reprogramming during the transition from monocytes to dendritic cells. (B) Genes linked to canonical DCs differentiation pathways. Significance thresholds were set at absolute LFC ≥ 0.5 and padj < 0.05. SLE: Systemic Lupus Erythematosus; Ctrl: healthy control; mo: monocytes; moDCs: monocyte-derived dendritic cells; and tolDCs: tolerogenic dendritic cells; DEGs: differential gene expression; padj: adjusted p-value; LFC: log₂FoldChange.

In the moDCs differentiation trajectory (Figure 4A), we found that *FADS2* was downregulated in SLE patients but upregulated in all other comparisons, suggesting impaired desaturation of polyunsaturated fatty acids in SLE (65). We also identified DEGs related to peroxisome proliferator-activated receptors (PPARs), transcription factors that regulate lipid metabolism and immune function, which have been reported to be significantly upregulated *in vitro* in moDCs generated using GM-CSF and IL-4 (65,66). Regarding tolDCs, differentiation and lipid metabolism, we observed downregulation of *CYP4F2* on both controls and SLE, consistent with reduced production of lipid-derived inflammatory mediators in tolDCs; previous studies have linked CYP4F2 lipid metabolism function with immune regulation (67). We also identified the downregulation of *SLC27A3* in tolDCs from controls and SLE (Figure 4A). Together, these findings underscore the importance of lipid metabolic reprogramming in shaping DCs differentiation and function (Figure 4A).

To further characterize differentiation trajectories, we analyzed genes annotated in the Gene Ontology Biological Process (GO:BP) category for DC differentiation. This exploration revealed additional transcriptional programs that define the functional identities of moDCs and tolDCs (68). In moDCs, *CCR6* was downregulated in both SLE and control samples, unlike in tolDCs, where its expression was preserved—consistent with a role for *CCR6* in supporting the migratory and regulatory features of the tolerogenic phenotype. Conversely, *CLEC4A* exhibited divergent regulation, being upregulated in moDCs from both SLE patients and healthy controls (Figure 4B). In tolDCs, *MERTK* was selectively upregulated in tolDCs from both SLE and controls; whereas STAT1 showed downregulation only in SLE.

Genes that displayed differential expression in only one of the differentiation trajectories (monocytes to moDCs or monocytes to tolDCs) within either SLE or control samples were considered trajectory-specific. Several genes showed exclusive or divergent patterns across groups. For example, *CCL5* and *PRTN3* were upregulated during monocyte-to-moDCs differentiation in healthy controls but were downregulated in all other trajectories, underscoring their context-dependent roles. *LAG3* also exhibited a disease-linked pattern, being downregulated in both SLE moDCs and tolDCs while remaining unchanged in controls (Figure 4B).

TolDCs can be generated through various stimuli, such as IL-10 or dexamethasone, each producing a distinct transcriptional profile (45). To assess how our differentiation conditions influenced the resulting transcriptomes, we compared the monocyte to tolDCs trajectory DEGs in both SLE and control samples with these previously reported stimulus-specific profiles, as well as with the general tolDCs signature described in the same study (45). In this analysis, we identified 71 DEGs associated with the tolDCs signature, of which 32 were previously identified as repressed and 41 as activated in tolDCs (45). This analysis of tolDCs signatures revealed both concordant and divergent gene expression patterns in our dataset. Several genes matched previously reported activation (13 genes) and repression profiles, while others showed opposite regulation (27 genes) (Supplementary Table S11). Interestingly, *IL7* displayed a disease-specific regulatory signature in SLE, being overexpressed in tolDCs from SLE patients but downregulated in tolDCs from controls, highlighting a potential disease-specific modulation of the tolerogenic signature. These differences appear to be influenced by variations in monocyte expression between patients and controls (Figure 5).

**Figure 5.**
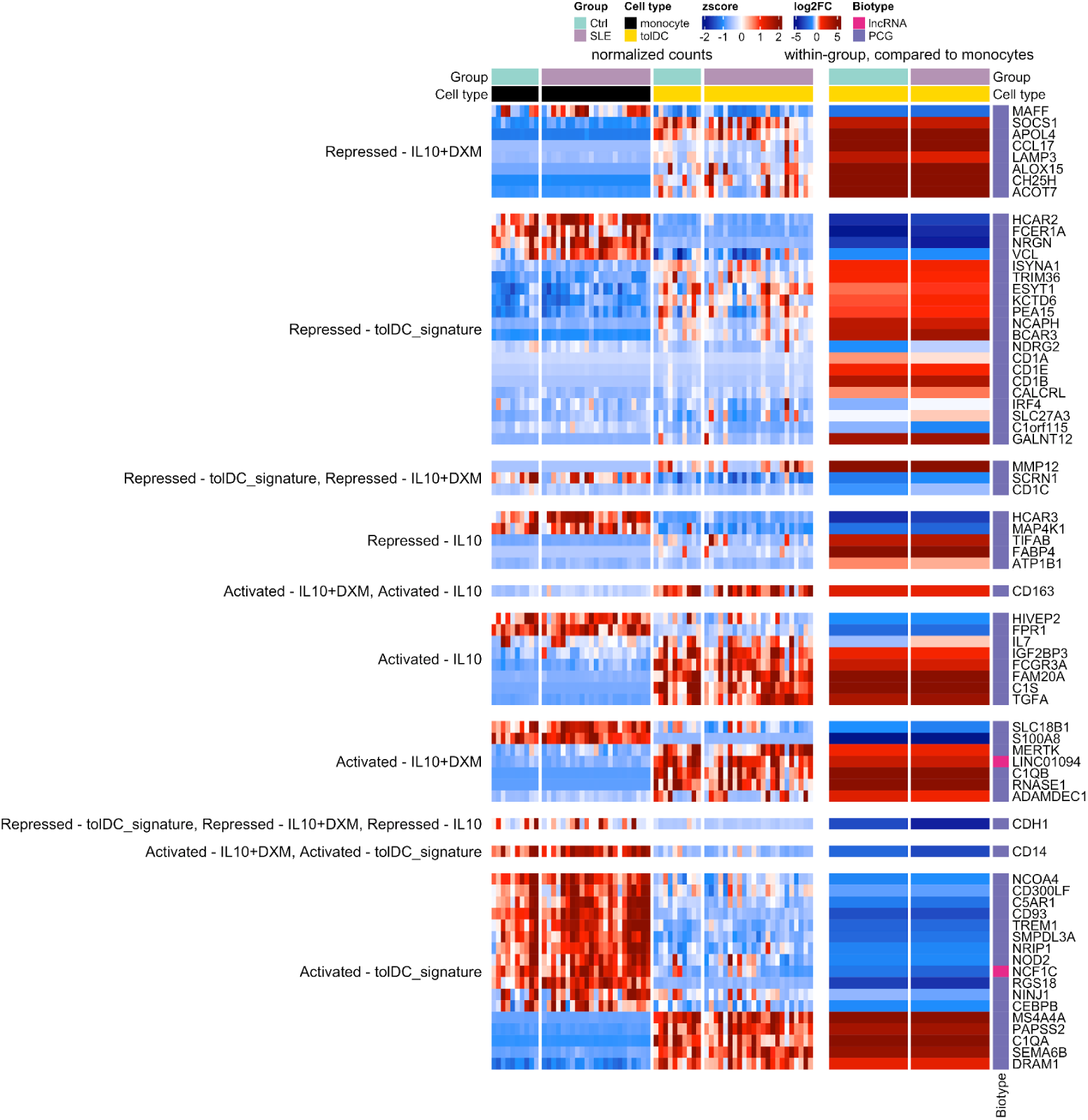
Transcriptional signatures associated with tolDCs obtained by SLE vs Controls comparison. Heatmaps showing 71 DEGs associated with tolDCs transcriptional signatures (32 downregulated and 41 upregulated) comparing monocytes vs DCs in SLE and controls. Rows represent genes and columns represent samples, colored according to cell types (monocytes, moDCs and tolDCs) and SLE vs Ctrl. Left panel: heatmap displays normalized gene expression values (z-score). Right panel: heatmap shows LFC values relative to monocytes.

Significance thresholds were set at absolute LFC ≥ 0.5 and padj < 0.05. Biotypes are color-coded: purple for protein-coding genes (PCG) and pink for long non-coding RNAs (lncRNAs). SLE: Systemic Lupus Erythematosus; Ctrl: healthy control; and tolDCs: tolerogenic dendritic cells; LFC: log₂FoldChange.

### Interaction of SLE on Monocyte to moDCs & tolDCs Transcriptional Programs

Our previous differentiation analysis provided a useful baseline for understanding how monocytes progress toward moDCs and tolDCs states in SLE and controls. However, this approach does not reveal whether the differentiation process unfolds differently in patients compared with controls. We therefore carried out a second analysis focused explicitly on identifying disease-dependent modifications to the differentiation trajectory, integrating SLE as an interaction factor (see Methods). We identified 33 DEGs exclusively in SLE (3 upregulated and 30 downregulated). Among these, 9 downregulated genes (*CCL2, EIF2AK2, HELZ2, HSPA1B, IFI6, ISG15, SIGLEC1, USP18,* and *XAF1*) were shared between moDCs and tolDCs (Figure 6A), mainly involved in interferon signaling, antiviral response, and immune regulation (Figure 6B). Seven downregulated genes specific to moDCs (*AGRN, IFITM1, OAS2, OAS3, RSAD2, SELL*, and *WIPF1*) (Figure 6A), were linked to cell adhesion, viral restriction, and innate immune activation (Figure 6B). Fourteen downregulated genes specific to tolDCs (*BST2, CYP51A1, DHX58, EBP, HERC5, IDI1, IFI35, IFIT3, IRF7, MSMO1, MUC1, MVD*, and *STARD4*) (Figure 6A), were associated with cholesterol biosynthesis, interferon-stimulated pathways, and transcriptional regulation of immune responses (Figure 6B). In contrast, two genes upregulated in tolDCs (*ABCG1* and *FTO*) were related to lipid transport and metabolic regulation, while *KDM1A/LSD1* upregulated in moDCs was linked to epigenetic modulation of transcription (Figure 6A; Figure 6B). These findings highlight that the transcriptional changes observed in DCs are strongly influenced by disease status, underscoring the weight of SLE in shaping the regulatory programs of monocyte-derived DCs.

**Figure 6.**
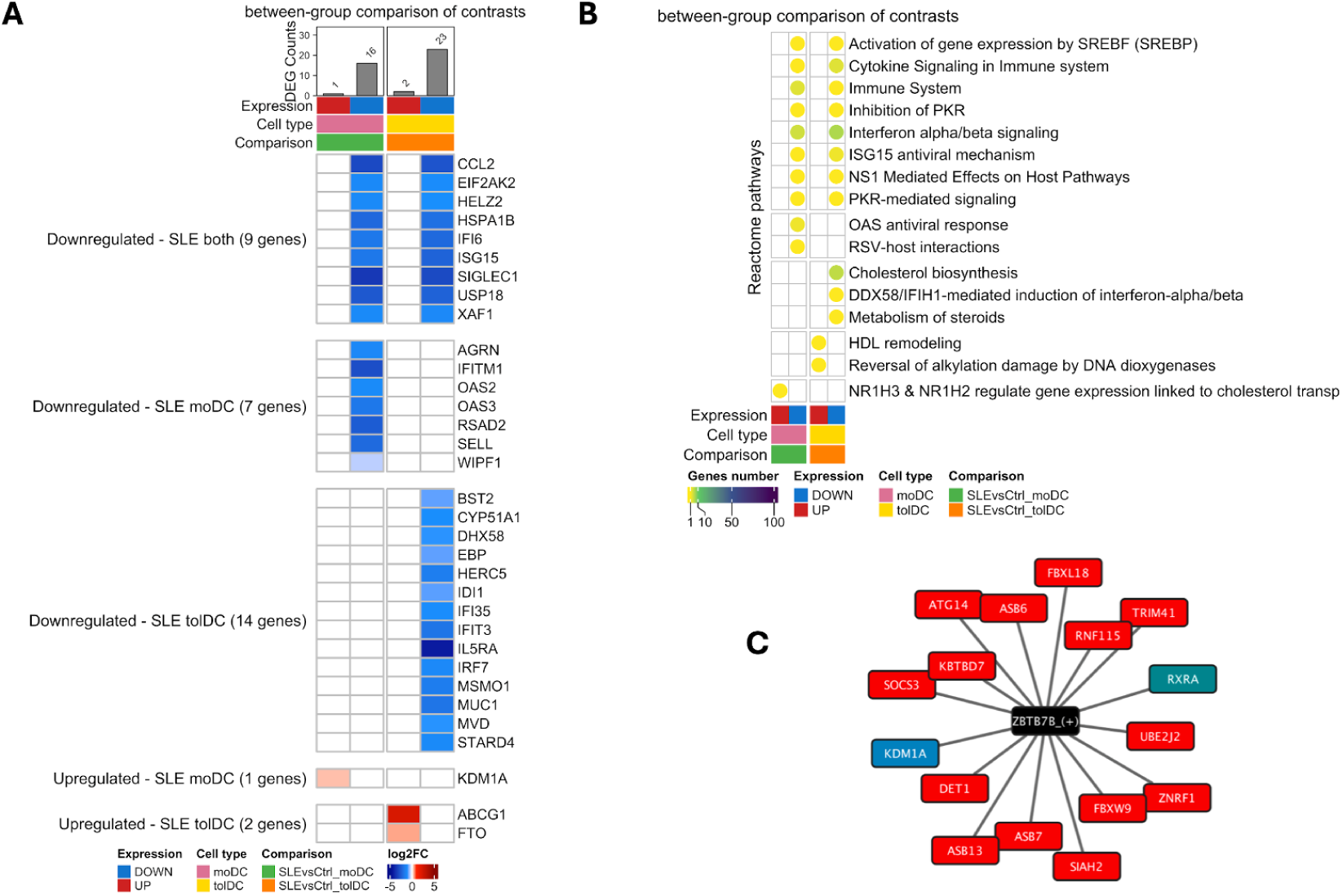
Monocyte-derived dendritic cells (moDCs and tolDCs) and associated transcriptional programs obtained by SLE vs. Ctrl comparison. (**A)** Heatmap of differentially expressed genes (DEGs) that are shared or unique to moDCs and tolDCs, using monocytes as baseline. **(B)** Significant biological processes implicated in SLE patients within these comparisons. (**C)** Network of *ZBTB7B* highlighted by pathway. Red: Targets associated with the global and biggest pathway (Class I MHC mediated antigen processing & presentation), Blue: *KDM1A*,/*LSD1* target from DE results associated with cholesterol transport and efflux, Green: *RXRA,* target associated with *KDM1A*/*LSD1* lipid metabolism.

To further investigate the transcriptional activity associated with this gene set, we constructed a targeted Gene Regulatory Network. Given that we did not find differential regulons between the SLE and control groups when introducing SLE interaction (Supplementary Figure S9; Supplementary Figure S10) and the PCA did not reveal global changes in regulon activity between SLE and controls (Supplementary Figure S11), we selected all regulons with gene targets among the DEGs shared or unique in moDCs and tolDCs (Supplementary Table S17), and focused on the regulon with the strongest differential activity: *ZBTB7B* (see Methods; Supplementary Figure S12). Notably, *ZBTB7B* directly targeted only one gene within the DEG set, *KDM1A/LSD1*, yet its regulon showed broad enrichment for antigen processing and fatty acid metabolic pathways when expanded to include additional *ZBTB7B* targets (Figure 6C).

Finally, we asked whether interferon-driven transcriptional programs present at baseline in monocytes are carried forward during differentiation into DCs. Focusing on genes shared between the monocyte SLE vs. control comparison and the interaction effect, we found that 18 of the 37 monocyte DEGs overlapped, including fourteen interferon-related transcription factors and their targets. These results indicate that type I interferon signalling imprinted in SLE monocytes is propagated into subsequent DC differentiation (Supplementary Table S18).

## Discussion

The breakdown of immune tolerance is a central mechanism of SLE pathogenesis, promoting the activation of autoreactive cells and the production of autoantibodies. Due to their pivotal role at the interface of innate and adaptive immunity, DCs are key in modulating the immune response, and their dysregulation contributes to both the breakdown of immune tolerance and chronic inflammation in SLE. A deeper understanding of the mechanisms underlying DCs functional phenotypes in people with SLE could provide significant insights for therapeutic approaches.

In this study, we compared the transcriptomic profiles of DCs derived from monocytes of Mexican women with SLE to those from women without SLE. We observed that ISGs constitute strong molecular markers across all three cell types, highlighting the relevance of interferon signaling in the pathophysiology of SLE, with multiple reports demonstrating that increased ISG expression parallels lupus initiation and its correlation with disease severity (69). We identified a substantial number of ISGs that were differentially expressed across the three cell types, related to interferon alpha/beta signaling, and other interferon-driven transcriptional pathways. Besides, monocytes showed the strongest enrichment of ISGs upregulation, exceeding the changes observed in DCs, such as *ISG15*, a negative regulator of type I IFN and previously reported to be upregulated in monocytes from SLE patients (69). This finding is consistent with the well-established role of type I IFN pathways in the pathogenesis of SLE, and it was related to the previously reported SLERRA signature (55). Suggesting that monocytes may contribute importantly to the SLE-associated interferon transcriptional profile, although the signature remains in derived DCs.

In addition to the enrichment of ISGs in monocytes, we identified several SLE biomarkers previously reported to be related to hub ISGs (62,70), including *IFI27, CMPK2, EIF2AK2, IFI44L, IFI6, IFIT1, ISG15, OAS1, OAS2, RSAD2, USP18,* and *LBH* (62,70). These findings are consistent with previous studies demonstrating that interferon signaling is central to the SLE pathogenesis: the overexpression of ISGs in blood, commonly known as “interferon signature”, is observed in over 80% of people with SLE and has been linked with disease activity (70–73). Among these, we detected increased expression of *IFI44L, IFI27, USP18* and *IFI6*, a set of genes previously proposed as a 4-gene model with strong diagnostic performance for interferonopathies (74). The presence of these genes as central nodes underscores their pivotal roles in interferon-driven transcriptional programs and in immune dysregulation in SLE.

In particular, *USP18* showed one of the highest fold changes (LFC = 3.123), highlighting its potential relevance to SLE pathophysiology given its role as a negative regulator of interferon signalling (75). Meanwhile, *IFI27* was overexpressed across the three cell types, underscoring its crucial role in its pathophysiology, likely reflecting its function in immune modulation, apoptosis regulation, and antiviral response (55,62). Importantly, interferon-inducible protein 44-like (*IFI44L*), has been shown to enhance the maturation of moDCs, upregulating key costimulatory molecules (e.g. CD40, CD80, CD83, and CD86) and promoting Th1/Th17 polarization in T cells, which may contribute to the breakdown of tolerance and chronic inflammation typical of SLE (76). Together, the persistent overexpression and functional contributions of these ISGs highlight their potential as therapeutic targets aimed at modulating IFN signaling in SLE.

Beyond the interferon-driven transcriptional signature, tolDCs from SLE patients exhibited changes enriched in metabolic and proliferative pathways. Genes involved in cholesterol and lipid metabolism (e.g. *ACAT2, MSMO1,* and *SMPD3*), lipid/cholesterol efflux (e.g. *ABCA1, ABCG1*), and regulators of the cell cycle (e.g. *CDCA5, CENPM*) were exclusively dysregulated in SLE tolDCs. These findings align with growing evidence regarding the role of lipid metabolism in the regulation of DCs responses; efficient cholesterol efflux and lipid metabolism is key to maintaining immune tolerance in DCs: impaired mechanisms promote accumulation of intracellular lipids, activate inflammasome and secrete pro-inflammatory cytokines, promoting autoreactive T and B cell responses and contributing to chronic inflammation and autoimmune responses (77–80). This dysregulation tolDCs from SLE, supports a model in which metabolic reprogramming contributes to DCs dysfunction, and highlights lipid homeostasis inducing strategies as potential targets to restore immune tolerance in SLE.

To further investigate differences between tolDCs and moDCs, we analyzed the differentiation trajectories from monocytes to each DC subtype and compared the resulting gene expression changes between SLE and controls. As expected, differentiation induced strong remodeling, with over 8,000 DEGs, including downregulation of monocyte markers (e.g. *CD14*), upregulation of DC maturation and antigen presentation markers (e.g. *CD40, CD80, CD68, HLA-DPA1,* and *HLA-DQA1*), anti-inflammatory mediators (e.g. *IL10, IL1R1*), and the downregulation of other classical maturation markers such as *CD83* (81). These transcriptional changes validate that our *in vitro* differentiation protocol successfully induces DCs features, providing a robust model to explore disease-driven alterations.

Beyond these expected changes, we identified shared transcriptional profiles of DCs differentiation in both SLE and controls (Figure 3). In moDCs, downregulation of *CXCL3, JAK3,* and *MAP3K8* suggests attenuation of early inflammatory signaling, while upregulation of *ADORA1/2B, CD1A, LIPA, TRAF3, CCR5, TNFRSF4, TSLP,* and *IRF4* reflects activation of pathways that support DCs maturation, lipid antigen handling, and purinergic regulation (82–86). Moreover, we identified lower expression of *IL10RA* in moDCs from SLE patients. Given that IL-10 signaling is key for preventing DCs activation and to induce differentiation towards tolerogenic phenotype through the IL10RA–JAK1–STAT3 axis, lower *IL10RA* expression may suggest a decreased ability of SLE moDCs to sense and respond to IL-10, despite the elevated IL-10 levels commonly observed in SLE (87,88). In contrast, moDCs from controls exhibited lower *IRF4* expression, a key driver of DCs maturation and antigen-presenting function; its reduced expression in controls is consistent with a physiologically restrained, steady-state maturation program (89–91). This higher *IRF4* gene expression in SLE moDCs likely reflects chronic *in vivo* inflammatory priming, consistent with the pre-activated status previously described in SLE myeloid cells (92,93). Together, the diminished *IL10RA* and dysregulated *IRF4* in SLE support the presence of cell-intrinsic defects in peripheral tolerance that may predispose antigen-presenting cells toward heightened activation.

Regarding tolDCs, both SLE and controls exhibited a transcriptional profile that resembles the human IL-10-producing DCs (DC-10) subset, characterized by high *IL10* and *CD163* expression, suggesting that IL-10–mediated differentiation efficiently drives these cells toward an immunosuppressive phenotype (87,94,95). Interestingly, despite this tolerogenic signature, we observed an unexpected upregulation of *HLA-DPA1* and *HLA-DQA*, which may reflect the preservation of antigen-presenting capacity; which under conditions of high IL-10 and reduced co-stimulation could support presentation of self-antigens with the aim of promoting T-cell anergy or regulatory T-cell induction (96,97). IL-4, one of the stimuli used in our differentiation protocol, further reinforces this phenotype by inducing sustained PPAR-γ activity, a pathway known to prevent full DCs maturation and limit pro-inflammatory T-cell priming (98). tolDCs displayed a coordinated transcriptional program characterized by elevated HLA class II expression together with IL-10/PPAR-γ–mediated immunoregulatory signaling, consistent with a phenotype that preserves antigen presentation within an anti-inflammatory context. Notably, tolDCs from SLE donors showed greater upregulation of the purinergic receptor ADORA2B, a regulator known to enhance tolerogenic reprogramming by promoting IL-10 production and suppressing pro-inflammatory cytokines (99,100). Although both SLE and controls exhibited a DC-10-like transcriptional profile, these results suggest different molecular pathways towards tolerogenic programming.

We next examined molecular pathways underlying the differences observed in the trajectory analysis comparing monocytes to moDCs and tolDCs in SLE and controls (Figure 4). This analysis revealed pronounced alterations in lipid metabolism, with several DEGs involved in lipid metabolism alongside increased expression of lipid-regulatory transcription factors such as PPARγ and PPARβ/δ (101,102). An interesting finding was the reduced expression of FADS2 and FADS1 in moDCs from SLE which are key enzymes required for generating long-chain polyunsaturated fatty acids and their pro-resolving derivatives (103). This downregulation aligns with the known defect in pro-resolving lipid mediator production in SLE and suggests impaired metabolic remodeling during DCs differentiation (104). These alterations may compromise the functional and immunoregulatory capacity of SLE moDCs.

Regarding other molecular pathways, we found that moDCs from controls exhibited the upregulation of *CCL5*, suggesting an enhanced capacity for chemotactic signaling and T cell recruitment, processes expected for DCs maturation (105). Additionally, we observed downregulation of *TMEM176B* and *TMEM176A*, genes previously associated with immature or regulatory states of DCs (106). These findings contribute to the idea that moDCs from controls can differentiate more effectively into an immunogenic phenotype than SLE moDCs (107–109).

In tolDCs from controls, the modest downregulation of BATF2, a transcription factor associated with IFN-mediated activation, might contribute to maintaining a moderate activation state that enables the rapid activation of tolerance mechanisms (110). BATF2 can interact with IRF family members, particularly IRF4, to shape enhancer activity and lineage-defining transcriptional programs; reduced BATF2 levels may limit pro-inflammatory BATF2–IRF4 complexes and instead favor IRF4-driven tolerogenic or homeostatic pathways (110). Consistent with this regulatory landscape, we also observed a mild upregulation of *STAT1* in control tolDCs compared with tolDCs from SLE patients. Given that STAT1 is essential for DCs differentiation, maturation, and IFN-responsive regulatory circuits, its higher expression in controls suggests preserved IFN-driven homeostatic signaling, supporting balanced antigen presentation and immune regulation (111,112). Interestingly, this regulatory balance appears altered in SLE, where both moDCs and tolDCs from SLE patients showed downregulation of *LAG3*, which could impair their capacity to deliver inhibitory signals to T cells and contribute to defective peripheral tolerance (113,114). Together, these pathways indicate that SLE monocytes carry a priming imprint that biases DCs differentiation toward a partially activated and metabolically dysregulated phenotype.

To situate our findings within an established framework, we adopted the consensus “tolDC signature” defined by Robertson *et al*. based on a comprehensive transcriptomic meta-analysis of diverse tolerogenic differentiation protocols. Using this reference allowed for systematic comparison between our tolDCs and a well-validated signature. In general, we observed strong replication of the reported expression profiles (Figure 5) (45), with the exception of IL-7, which was upregulated only in SLE tolDCs. This selective upregulation may reflect the chronic inflammatory nature of SLE, and could contribute to DCs maturation and pro-inflammatory signaling, consistent with IL-7’s broader role in supporting autoreactive responses in autoimmune diseases (115).

Our findings reveal that the transcriptional programs governing dendritic cell differentiation are profoundly reshaped in the context of SLE (116). By performing an interaction analysis, we identified disease-specific gene expression changes that distinguish moDCs and tolDCs from their healthy counterparts. The predominance of downregulated genes, particularly those involved in interferon signaling, antiviral responses, and immune regulation, underscores the extent to which SLE disrupts canonical pathways essential for DC function (117–122). The downregulation of negative regulators of IFN signaling in moDCs and tolDCs suggests a loss of feedback control mechanisms (123). This impaired regulation may perpetuate chronic interferon activity, thereby sustaining inflammatory programs and compromising tolerogenic differentiation in SLE DCs.

Gene regulatory networks built from the transcriptome analysis of the differentiation of monocytes to DCs in SLE and controls revealed a shared transcriptional architecture shaping DCs differentiation with a regulatory signature in SLE. In particular, SLE DCs exhibited a higher number of unique regulons, suggesting that SLE monocytes undergo DCs differentiation with a pre-existing regulatory bias, likely shaped by chronic inflammation and IFN signaling. One of the most prominent signals was the CREB1 regulon, upregulated during moDCs differentiation and known to promote DC survival, activation, and IL-10–driven immunomodulatory functions (124,125).

Our regulatory networks analysis highlighted the involvement of less-characterized zinc-finger/BTB transcription factors, particularly ZBTB7B, which was upregulated during moDCs differentiation and has recently been implicated in regulating myeloid commitment, differentiation, and maturation in murine models (126). Although classically associated with T-cell development, its significant activity in our human *in vitro* system suggests a potential role in shaping DCs development and homeostasis (126). Notably, ZBTB7B showed the largest regulon-level activity difference between SLE and controls, and was borderline enriched in the interaction analysis, pointing to a possible contribution to tolerance induction in the SLE context (127). While it directly regulates only one upregulated DEG in moDCs (KDM1A/LSD1; Figure 6), ZBTB7B appears to sit atop a broader regulatory module linking cholesterol transport, fatty-acid metabolism, and MHC-I antigen presentation, processes known to be dysregulated in SLE DCs (78). During tolDCs differentiation, additional regulators such as ZKSCAN1 and KLF16 were induced, whereas inflammatory drivers including RELA, BCL3, and IRF3 were downregulated (Figure 3), consistent with a coordinated program favoring a tolerogenic, non-inflammatory phenotype (128–134). Although these findings point to promising regulatory candidates involved in monocyte DCs differentiation, potentially altered in SLE, their roles remain correlative and require functional validation to confirm mechanistic relevance.

Although the transcriptional profile of our tolDCs resembles the well-described DC-10 phenotype, our study is limited by the absence of functional validation. Determining whether these *in vitro*-generated cells recapitulate the immunosuppressive properties of bona fide tolDCs will require additional work. Future studies should include flow cytometric assessment of canonical DC-10 markers (CD141, CD163, HLA-G, ILT4/ILT receptors), quantification of IL-10 production and suppression of pro-inflammatory cytokines, and co-culture assays with naïve CD4⁺ T cells to evaluate their ability to induce regulatory T cells. Incorporating these functional assays will be essential to establish whether the observed transcriptomic programs translate into the tolerogenic activity expected of tolDCs and to refine our understanding of how these pathways are altered in SLE.

## Conclusion

Our findings demonstrate an increased expression of interferon-stimulated genes in SLE patient monocytes relative to controls, thereby confirming a robust ISG signature that persists as these cells differentiate into DC subsets. Both moDCs and tolDCs exhibited disease specific transcriptional remodeling involving immune signaling pathways, lipid metabolism, and epigenetic regulators. The analysis of the tolerogenic signature revealed areas of concordance and divergence from canonical *in vitro* models, suggesting that tolerogenic programming is strongly shaped by the cellular and inflammatory context of SLE.

Although transcriptional differences in tolDCs between SLE and controls were generally subtle, the trajectory analyses indicate that key mechanisms required for establishing tolerogenic capacity may be disrupted in SLE. Together, these findings highlight the importance of characterizing tolDCs heterogeneity and their regulatory interactions more deeply, particularly in light of the regulons implicated here. Importantly, by clarifying how DCs differentiation is altered in SLE, this work provides a foundation for identifying molecular pathways that could be therapeutically targeted to reinforce tolerogenic DCs function and ultimately help restore immune tolerance in SLE.

## Data Availability Statement

Raw sequencing data (FASTQ and BAM files) generated in this study are publicly available in the European Nucleotide Archive (ENA) under accession number PRJEB104208 (release state: 26th Nov 2026). All analysis code, processed results, and figures are openly available in Zenodo at https://doi.org/10.5281/zenodo.17742206.

All results from the regulatory network analysis of SLE-associated regulon activity using the SCENIC pipeline are available in Zenodo at https://doi.org/10.5281/zenodo.17419882.

Additionally, a Quarto book containing the full codebase, documentation, and reproducible workflows is hosted on GitHub at https://github.com/NeuroGenomicsMX/SLE_Transcriptomic_analysis_tolerogenic.

Together, these repositories ensure open access, scientific transparency, and full reproducibility of the study.

## Author Contributions

ALHL contributed to conceptualization, data curation, formal analysis, funding acquisition, investigation, methodology development, resource provision, software implementation, validation, visualization, writing of the original draft, and manuscript review and editing. ELCN and SSM contributed to data curation, formal analysis, investigation, methodology development, software implementation, validation, visualization, and both original draft preparation and manuscript editing. DRE contributed to formal analysis, investigation, methodology, software, validation, visualization, and manuscript writing and editing. AP, LTN, ETV, and GFR contributed essential resources and manuscript review. GFV contributed to conceptualization, investigation, resource provision, and manuscript editing. JSGS contributed to formal analysis, project administration, supervision, validation, and manuscript editing. GT contributed to conceptualization, investigation, and manuscript editing. FR contributed to conceptualization, project administration, formal analysis, methodology, project administration, funding acquisition, supervision, and manuscript editing. SLFV contributed to conceptualization, methodology, supervision, project administration, funding acquisition, and manuscript editing. MGA contributed to conceptualization, methodology, supervision, funding acquisition, and manuscript editing. DAR contributed to conceptualization and manuscript editing. AMR contributed to conceptualization, funding acquisition, methodology, project administration, supervision, writing of the original draft, and writing review and editing.

## Acknowledgments

ALHL is a doctoral student from Programa de Doctorado en Ciencias Biomédicas, Universidad Nacional Autónoma de México (UNAM) and has received fellowship CVU (No. [711015]) from Secretaría de Ciencias, Humanidades, Tecnología e Innovación (SECIHTI, formerly CONAHCYT).

ELCN is a postdoctoral researcher at Laboratorio Internacional de Investigación sobre el Genoma Humano (LIIGH-UNAM) and she holds a fellowship from SECIHTI, formerly CONAHCYT (CVU 781634). ELCN acknowledges the use of Microsoft Copilot, a generative AI technology developed by Microsoft, for support in refining text. All content generated was reviewed and verified by the author for accuracy and compliance with Frontiers guidelines.

This work received support from Luis Aguilar, Alejandro León, and Jair García of the Laboratorio Nacional de Visualización Científica Avanzada. We also thank Carina Uribe Díaz, and Alejandra Castillo Carbajal, Karen Julia Nuñez-Reza and Christian Molina-Aguilar for their technical support. Authors would like to express their special acknowledgment to Fundación Proayuda Lupus Morelos A.C, LupusMX, El despertar de la Mariposa, Centro de Estudios Transdisciplinarios Athié-Calleja por los Derechos de las Personas con Lupus A. C. (Cetlu A.C.), Lupus en Yucatán, Asociación de Lupus y AIJ caminando Juntos A.C., Lupus Tuxtla, Aprendiendo a Vivir con Lupus y Fibromialgia A.C., Realidad Lúpica, AMMA Abraza con Amor A.C., for their invaluable support. We thank Mauricio Guzman for design and styling support.

## Abbreviations

SLE: Systemic Lupus Erythematosus
Ctrl: healthy control
mo: monocytes
moDCs: monocyte-derived dendritic cells
tolDCs: tolerogenic dendritic cells
DEGs: differential gene expression
padj: adjusted p-value
LFC: log₂FoldChange
FDR: false discovery rate
BH: Benjamini–Hochberg procedure
IFN: interferon
PBMC: peripheral blood mononuclear cell
PCA: principal component analysis

## Publisher’s Note

All claims expressed in this article are solely those of the authors and do not necessarily represent those of their affiliated organizations, or those of the publisher, the editors and the reviewers. Any product that may be evaluated in this article, or claim that may be made by its manufacturer, is not guaranteed or endorsed by the publisher.

