## Supplemental figures for "Gene expression profiling of dendritic cell tolerance dysfunction in women with Systemic lupus erythematosus"

### Supplementary information

**Table S1**. Sociodemographic and treatment data used for normality assessment and descriptive analyses, GCC: glucocorticoids, AM: antimalarials, MMF: mycophenolate mofetil, AZT: azathioprine, MTT: methotrexate, PDN: prednisone, PDS: prednisolone, Ctrl: non-SLE affected controls, SLE: Systemic lupus erythematosus.

**Table S2.** Information of monocyte, dendritic cells (tolDCs and moDCs) generated from systemic lupus erythematosus (SLE) patients and healthy controls (CSV format).

**Table S3.** Distribution and normality assessment of continuous variables.

**Table S4.** Table of metadata and sample IDs of all transcriptomes used in this project (CSV format).

**Table S5.** DEGs obtained by comparing SLE vs. Ctrl in each cell type (Differentiation comparison) (CSV format).

**Table S6.** Table of biological processes involved with the DEGs obtained by comparing SLE vs. Ctrl in each cell type (Differentiation comparison) (TSV format).

**Table S7.** Table of biological processes involved with SLE-associated biomarkers obtained by comparing SLE vs. Ctrl in each cell type (Differentiation comparison) (CSV format).

**Table S8.** Table of biological processes involved with the DEISGs obtained by comparing SLE vs. Ctrl in each cell type (Within-group comparison) (CSV format).

**Table S9.** DEGs obtained by comparing DC (moDC and tolDC) vs monocytes from SLE vs. Ctrl (Within-group comparisons)

**Table S10.** List of DEGs previously reported.

**Table S11.** List of GO terms associated to previously reported genes.

**Table S12.** Table of biological processes involved by comparing DC (moDC and tolDC) vs monocytes from SLE vs. Ctrl (Within-group comparisons)

**Table S13.** DEGs obtained by comparing DC (moDC and tolDC) vs monocytes from SLE vs. Ctrl, to capture **disease-specific modifications of monocyte-to-dendritic cell differentiation programs** (Interaction effect)

**Table S14.** Intersection of DEGs across differentiation comparison, within-group comparisons, and interaction effect

**Table S15.** Intersection of DEGs in monocytes from differentiation comparison and interaction effect (Differentiation comparison + Interaction effect)

**Table S16.** Regulon activity scores across samples, measured by Scenic's normalized AUC values expressed as z-scores.

**Table S17.** Regulon activity scores directly targeting DEGs, measured by Scenic's normalized AUC values expressed as z-scores.

**Table S18:** Shared monocyte gene expression patterns and Log Fold Changes from Differentiation (SLE vs. Control) and Interaction effect (reference: control monocytes, including moDCs and tolDCs).

#### Supplementary Figures

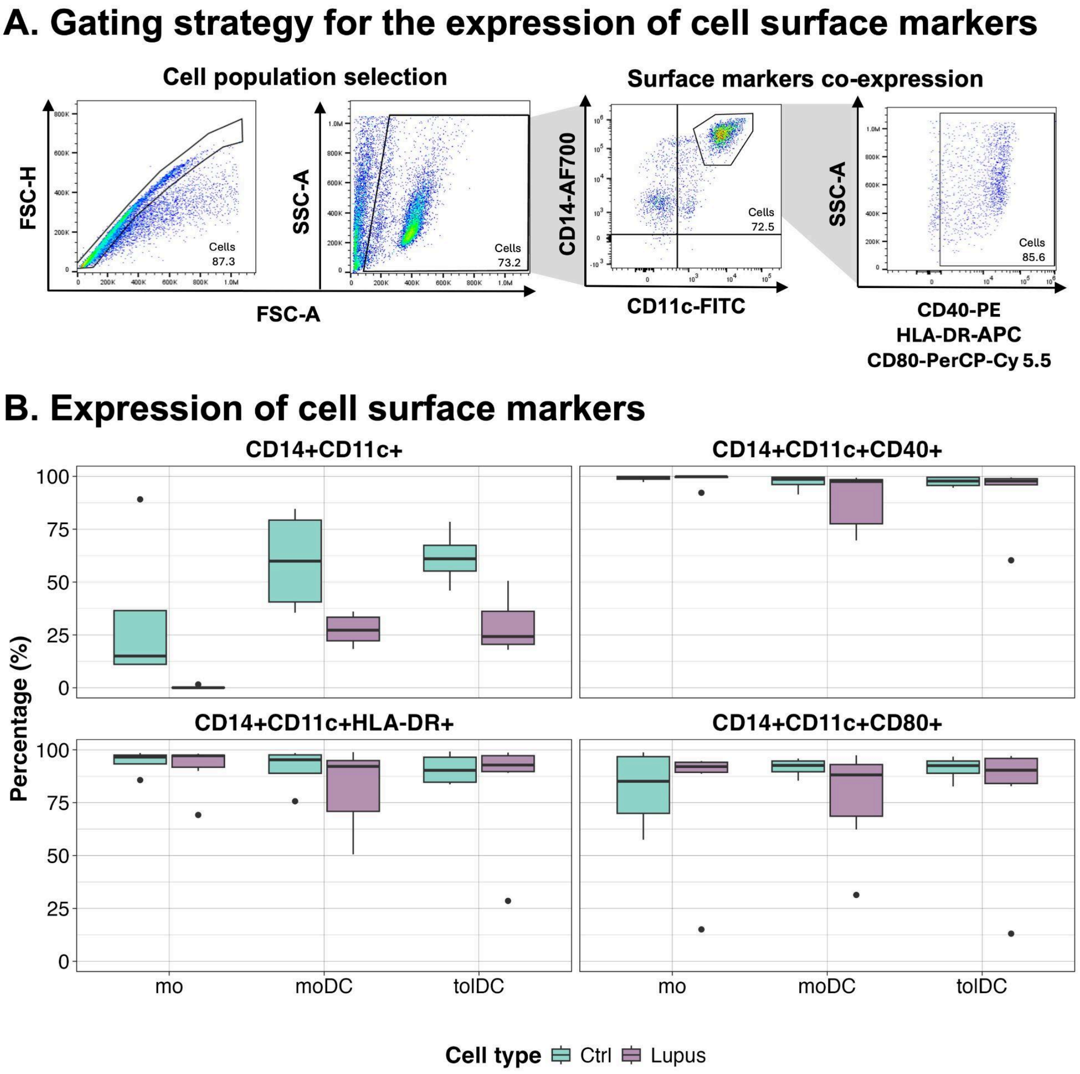


**Supplementary Figure S1. Flow cytometric analysis and quantitative bar chart analyses of surface marker expression. (A)** Gating strategy for the analysis of monocyte and dendritic cells surface marker expression. **(B)** Expression of CD14, CD11c, HLA-DR, CD80 and CD40 are shown among monocytes, monocyte-derived dendritic cells (moDCs), and tolerogenic dendritic cells (tolDCs) of people with SLE (purple bars) and controls (aqua bars).

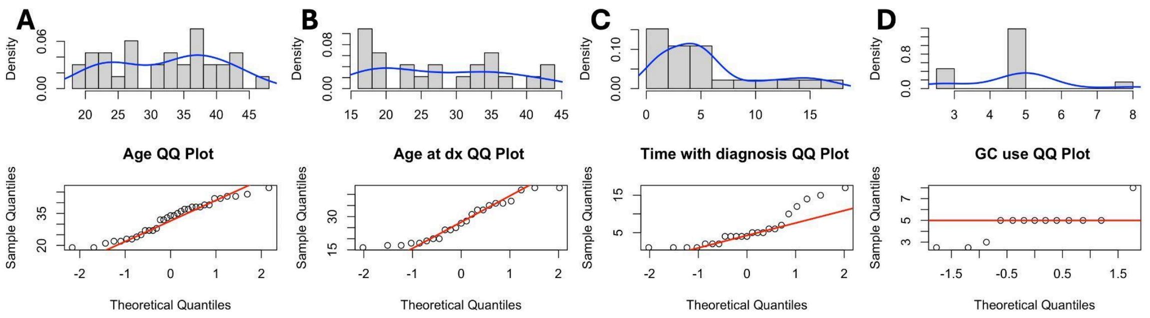


**Supplementary Figure S2. Distribution and normality assessment of continuous variables.** Barplots are divided into panels showing density distributions and Q–Q plots for **(A)** Age, **(B)** Age at treatment initiation, **(C)** Time since diagnosis, and **(D)** corticosteroid use **(QC).** Panel **(E)** displays the scatterplot of sample quantiles versus theoretical quantiles. These plots illustrate the distributional properties of the variables and their deviation from normality, as assessed by Shapiro–Wilk tests.

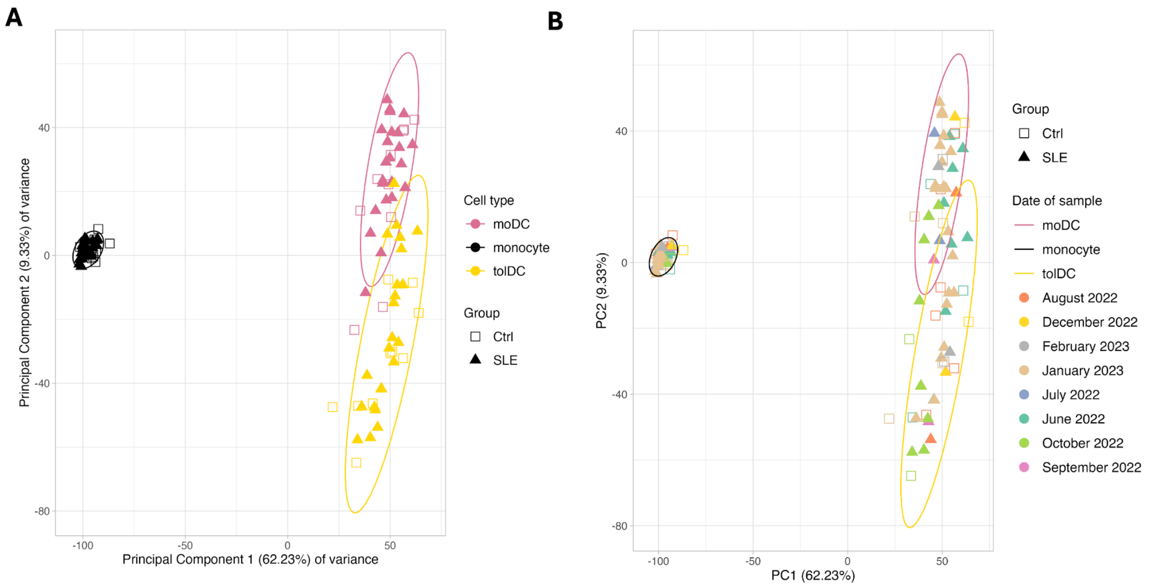


**Supplementary Figure S3. Principal Component Analysis (PCA) of normalized RNA-seq counts. A)** PCA colored by Cell type (monocytes, moDC, and tolDC) and shaped by Group (SLE and Ctrl), showing clear segregation primarily driven by cell identity. **B)** PCA plot including sampling dates (collection date) to evaluate potential batch effects across collections, revealing no major clustering by date. The first two principal components (PC1 and PC2) explain 62.23% and 9.33% of the total variance, respectively. Each point represents one sample. Abbreviations: SLE, systemic lupus erythematosus; Ctrl, healthy control; mo, monocytes; moDC, monocyte-derived dendritic cells; tolDC, tolerogenic dendritic cells. The data underlying this figure can be found in Supplementary Table S1 and Dataset S2.

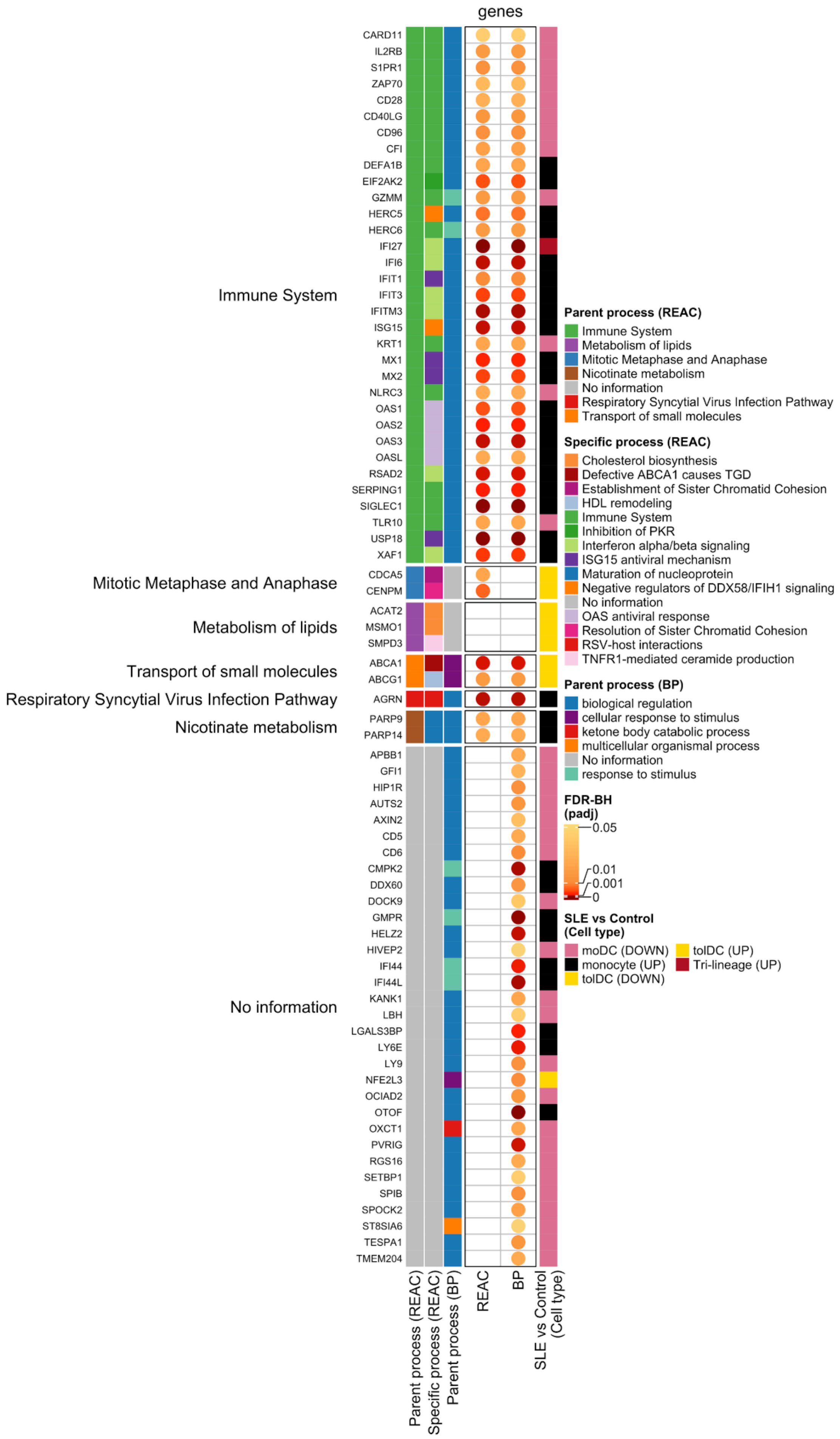
genes
Immune System
Mitotic Metaphase and Anaphase
Metabolism of lipids
Transport of small molecules
Respiratory Syncytial Virus Infection Pathway
Nicotinate metabolism
No information
CARD11
IL2RB
S1PR1
ZAP70
CD28
CD40LG
CD96
CFI
DEFA1B
EIF2AK2
GZMM
HERC5
HERC6
IFI27
IFI6
IFIT1
IFIT3
IFITM3
ISG15
KRT1
MX1
MX2
NLRC3
OAS1
OAS2
OAS3
OASL
RSAD2
SERPING1
SIGLEC1
TLR10
USP18
XAF1
CDCA5
CENPM
ACAT2
MSMO1
SMPD3
ABCA1
ABCG1
AGRN
PARP9
PARP14
APBB1
GFI1
HIP1R
AUTS2
AXIN2
CD5
CD6
CMPK2
DDX60
DOCK9
GMPR
HELZ2
HIVEP2
IFI44
IFI44L
KANK1
LBH
LGALS3BP
LY6E
LY9
NFE2L3
OCIAD2
OTOF
OXCT1
PVRIG
RGS16
SETBP1
SPIB
SPOCK2
ST8SIA6
TESPA1
TMEM204
Parent process (REAC)
Specific process (REAC)
Parent process (BP)
REAC
BP
SLE vs Control (Cell type)
Parent process (REAC)
Immune System
Metabolism of lipids
Mitotic Metaphase and Anaphase
Nicotinate metabolism
No information
Respiratory Syncytial Virus Infection Pathway
Transport of small molecules
Specific process (REAC)
Cholesterol biosynthesis
Defective ABCA1 causes TGD
Establishment of Sister Chromatid Cohesion
HDL remodeling
Immune System
Inhibition of PKR
Interferon alpha/beta signaling
ISG15 antiviral mechanism
Maturation of nucleoprotein
Negative regulators of DDX58/IFIH1 signaling
No information
OAS antiviral response
Resolution of Sister Chromatid Cohesion
RSV-host interactions
TNFR1-mediated ceramide production
Parent process (BP)
biological regulation
cellular response to stimulus
ketone body catabolic process
multicellular organismal process
No information
response to stimulus
FDR-BH (padj)
0.05
0.01
0.001
0
SLE vs Control (Cell type)
moDC (DOWN)
monocyte (UP)
tolDC (DOWN)
tolDC (UP)
Tri-lineage (UP)

**Supplementary Figure S4. Biological processes involved and associated with the DEGs obtained by comparing SLE vs. Ctrl in each cell type.** Processes were classified according to Reactome (REAC) parent and specific categories, and Gene Ontology Biological Process (GO:BP) parent terms. REAC parent processes include Immune system, Synthesis of phosphatidylinositol (PI), Cell division, Lipid metabolism and Transport of small molecules. Color scale represents padj (FDR–BH correction), with a maximum threshold of padj < 0.05. DEG: Differential expressed genes. The data underlying this figure can be found in Supplementary Table S6.

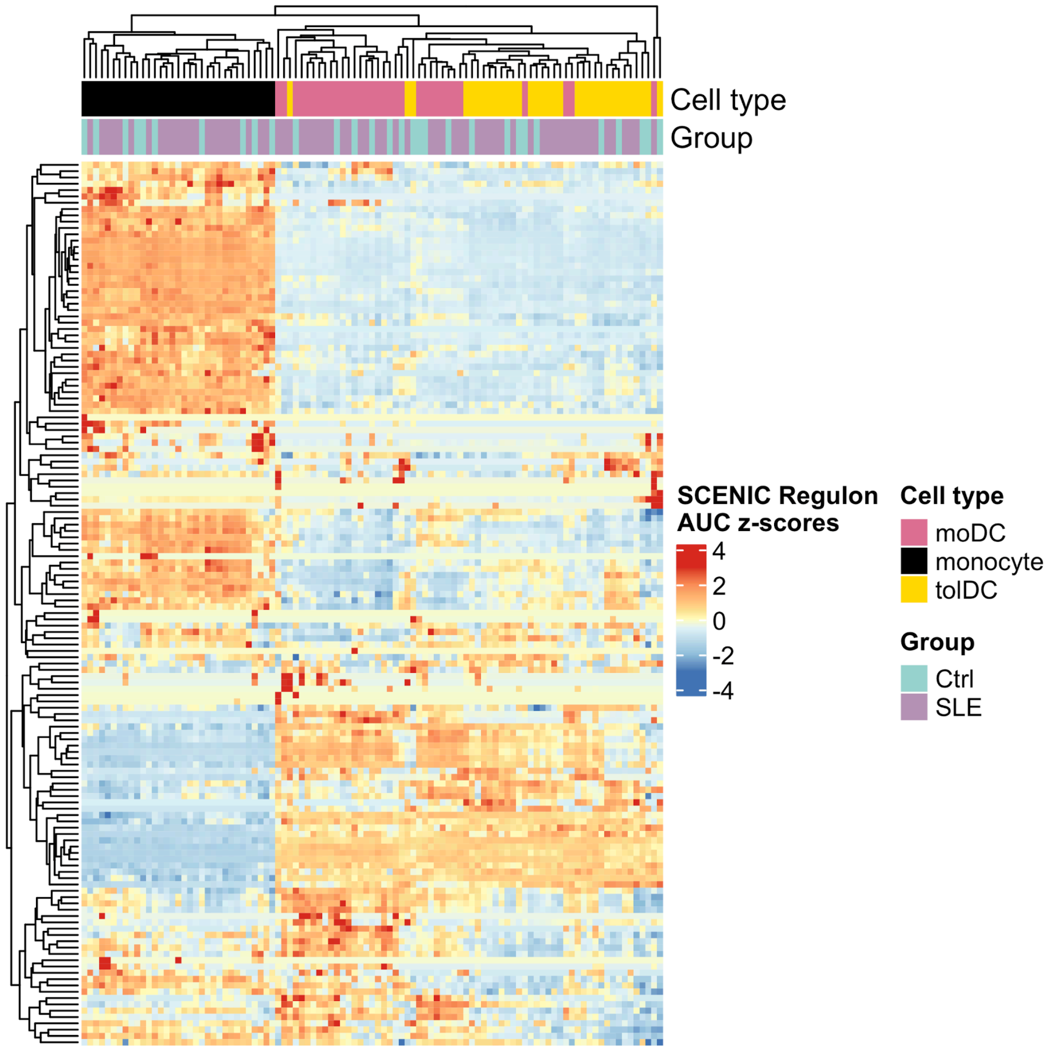


**Supplementary Figure S5. Regulon activity scores (AUC, z-score normalized).** Heatmap representation of regulon activity across samples, showing normalized AUC values expressed as z-scores. Rows correspond to individual regulons, while columns represent samples according to conditions (SLE or Ctrl). Color intensity reflects relative activity, highlighting the regulons with increased or decreased activity patterns. The data underlying this figure can be found in Supplementary Table S14.

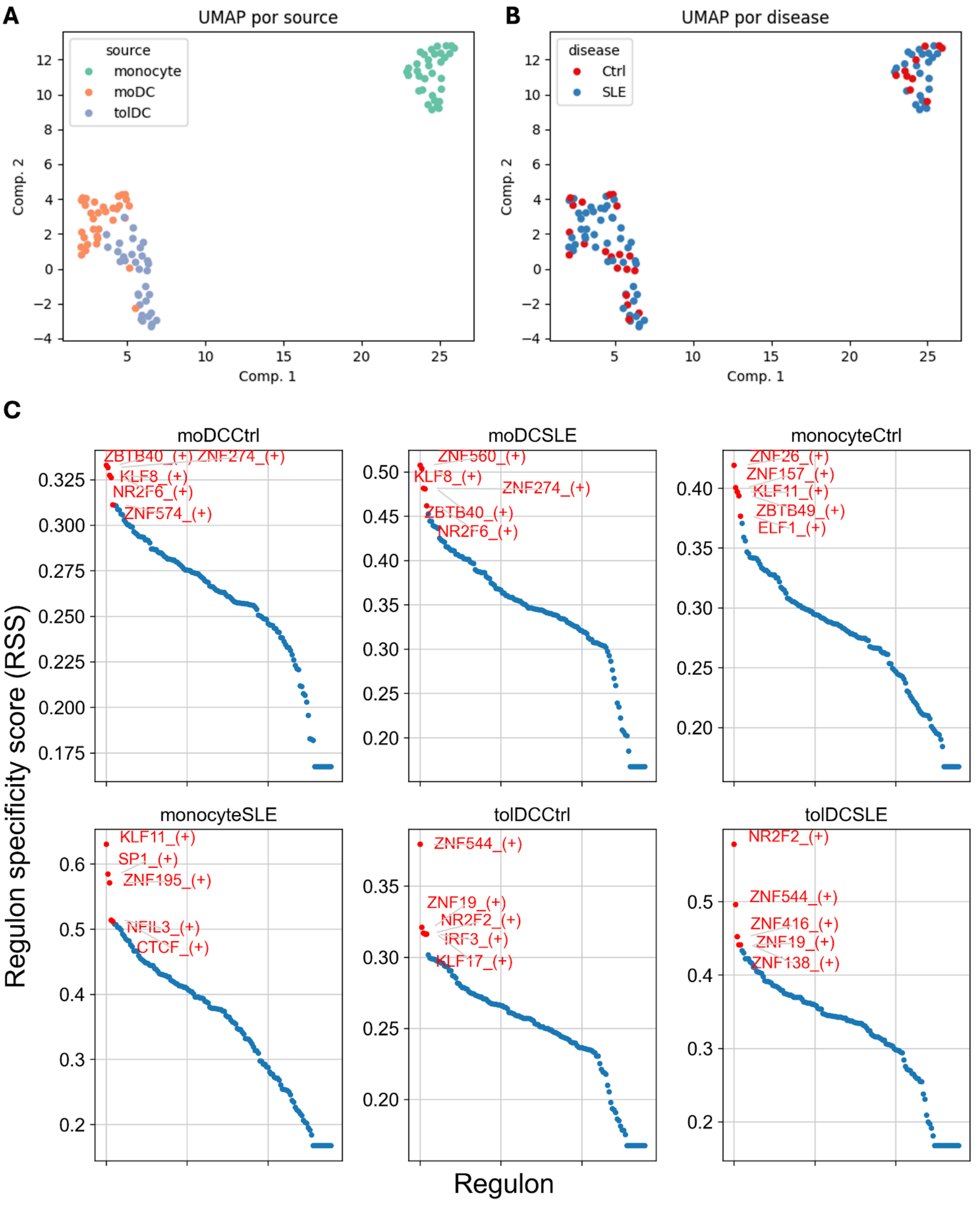


**Supplementary Figure S6. UMAP visualization and regulon specificity analysis of monocyte-derived populations. (A)** UMAP projection based on components 1 and 2, colored by cell type (monocytes in green, moDC in orange, and tolDC in blue), showing clear segregation of samples according to cell identity. **(B)** UMAP projection of the same dataset, colored by group (SLE in blue and Ctrl in red), illustrates the distribution of samples across disease conditions. **(C)** Regulon Specificity Score (RSS) analysis highlighting the top five transcription factors (TFs) identified with pySCENIC. The knee plot depicts the ranking of regulons by specificity score. The data underlying this figure were derived from the file *multi_runs_regulons_auc_trk.loom* and are available at Zenodo, DOI: 10.5281/zenodo.17419882.

Supplementary Figure S7

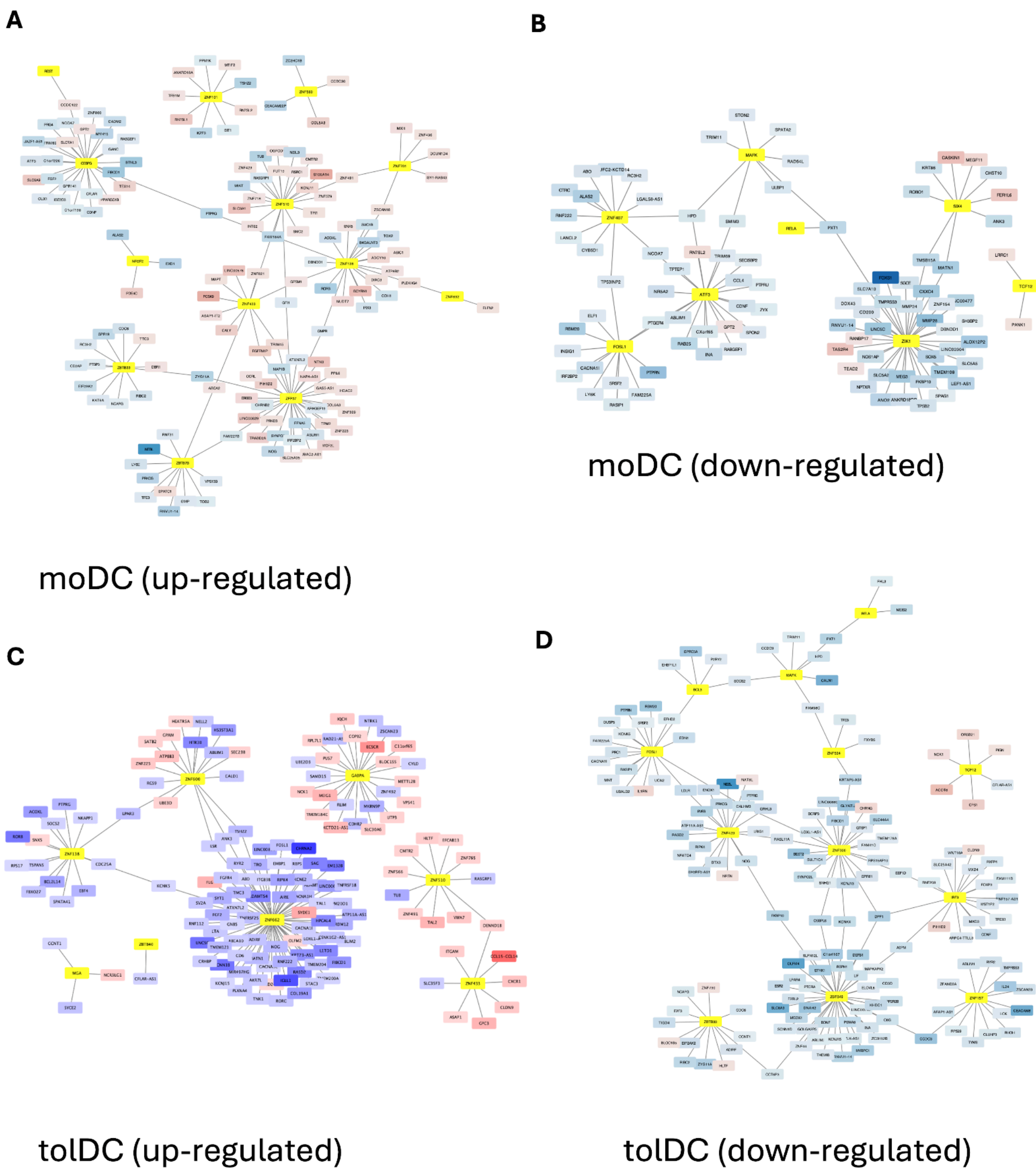


**Supplementary Figure S7. Regulatory networks exclusive to SLE during moDC and tolDC differentiation**. Depiction of regulons significantly **A)** upregulated and **B)** downregulated upon differentiation of moDC, **C)** upregulated and **D)** downregulated upon differentiation of tolDC, exclusive to SLE. Each node represents an identified regulon, with edges indicating inferred regulatory interactions. The contrast between the two networks highlights transcriptional programs activated and repressed in the SLE context, underscoring the disease-specific nature of these regulatory changes. SLE: Systemic Lupus Erythematosus; moDC: monocyte-derived dendritic cells.

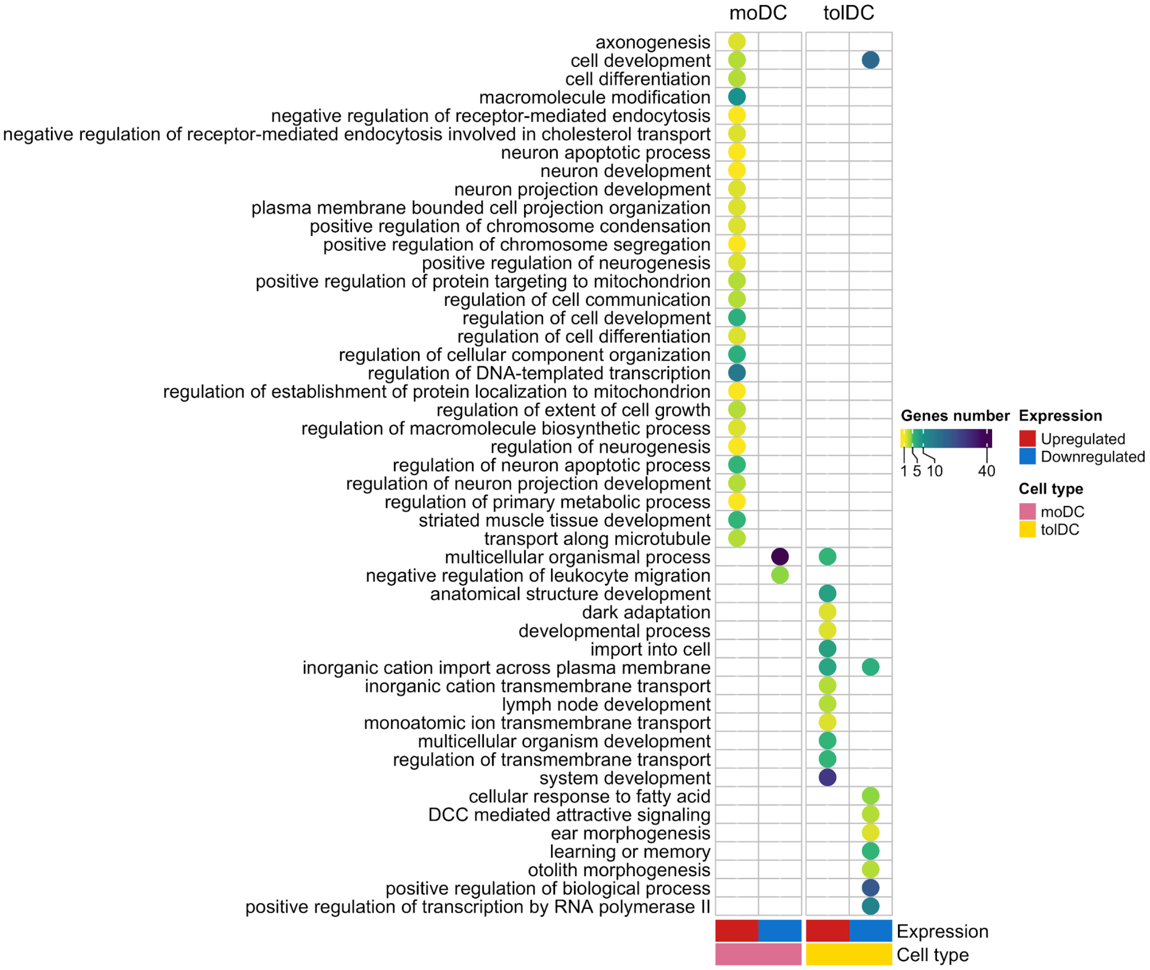


**Supplementary Figure S8. Gene enrichment analysis of SLE-exclusive regulons involved in the differentiation to moDC and tolDC.** Gene Ontology Biological Process (GO:BP) and Reactome (REAC) pathway terms enriched among the target genes of moDC and tolDC SLE-specific regulons. These target genes were also significantly differentially expressed during the differentiation process. Terms are listed in rows. Most terms correspond to GO:BP, with the exception of 'DCC mediated attractive signaling,' which is derived from the REAC database. Columns indicate the cell type associated with each regulon (annotated as "Cell type"). The "Expression" annotation denotes whether each regulon set was up or downregulated during differentiation to the corresponding cell type. The color scale represents the number of differentially expressed target genes associated with each term.

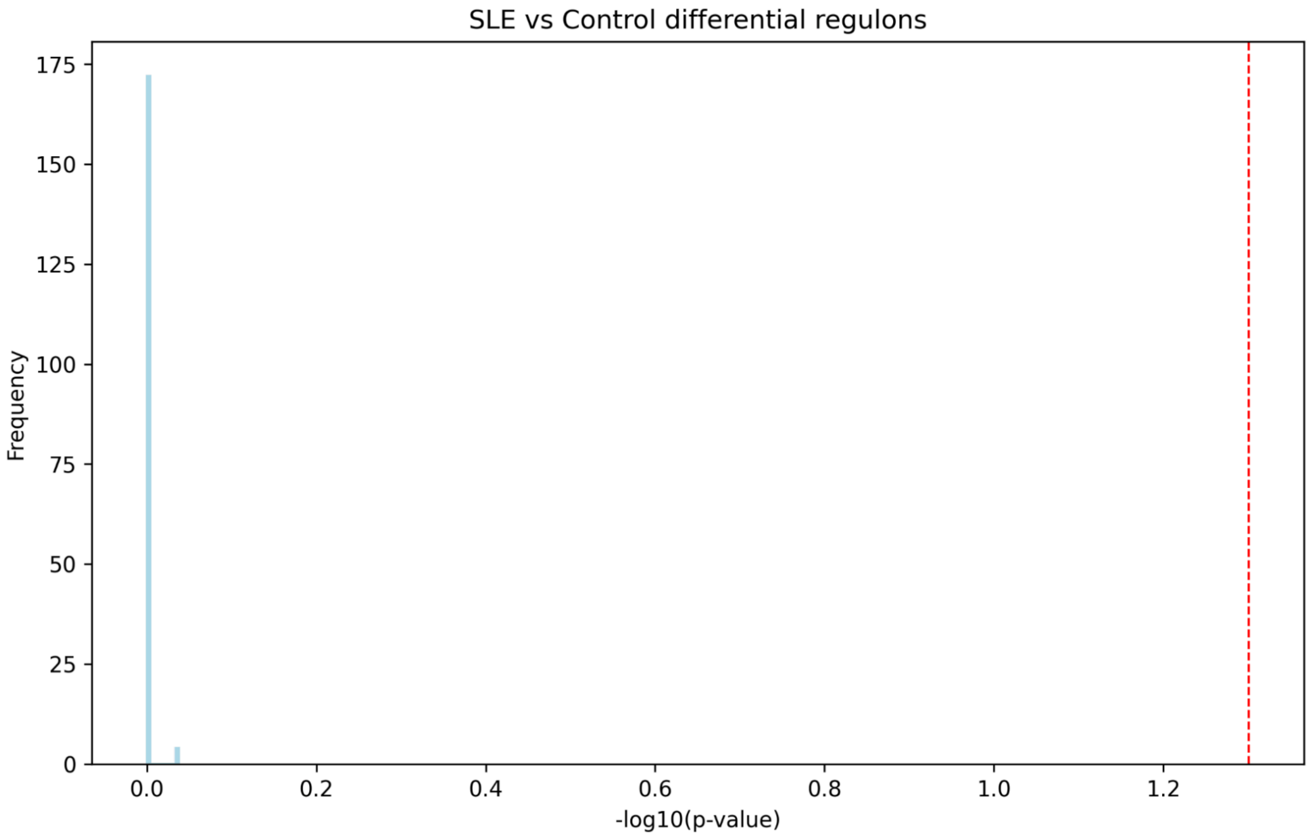


**Supplementary Figure S9.** Differential regulons adjusted p-values from the SLE vs control comparison Histogram of p-values from the differential analysis of regulons in SLE patients compared to controls. P-values were adjusted for multiple testing using the Benjamini-Hochberg procedure (FDR < 0.05) and -log10 transformed. The red line shows the significant cut-off threshold at 0.05.

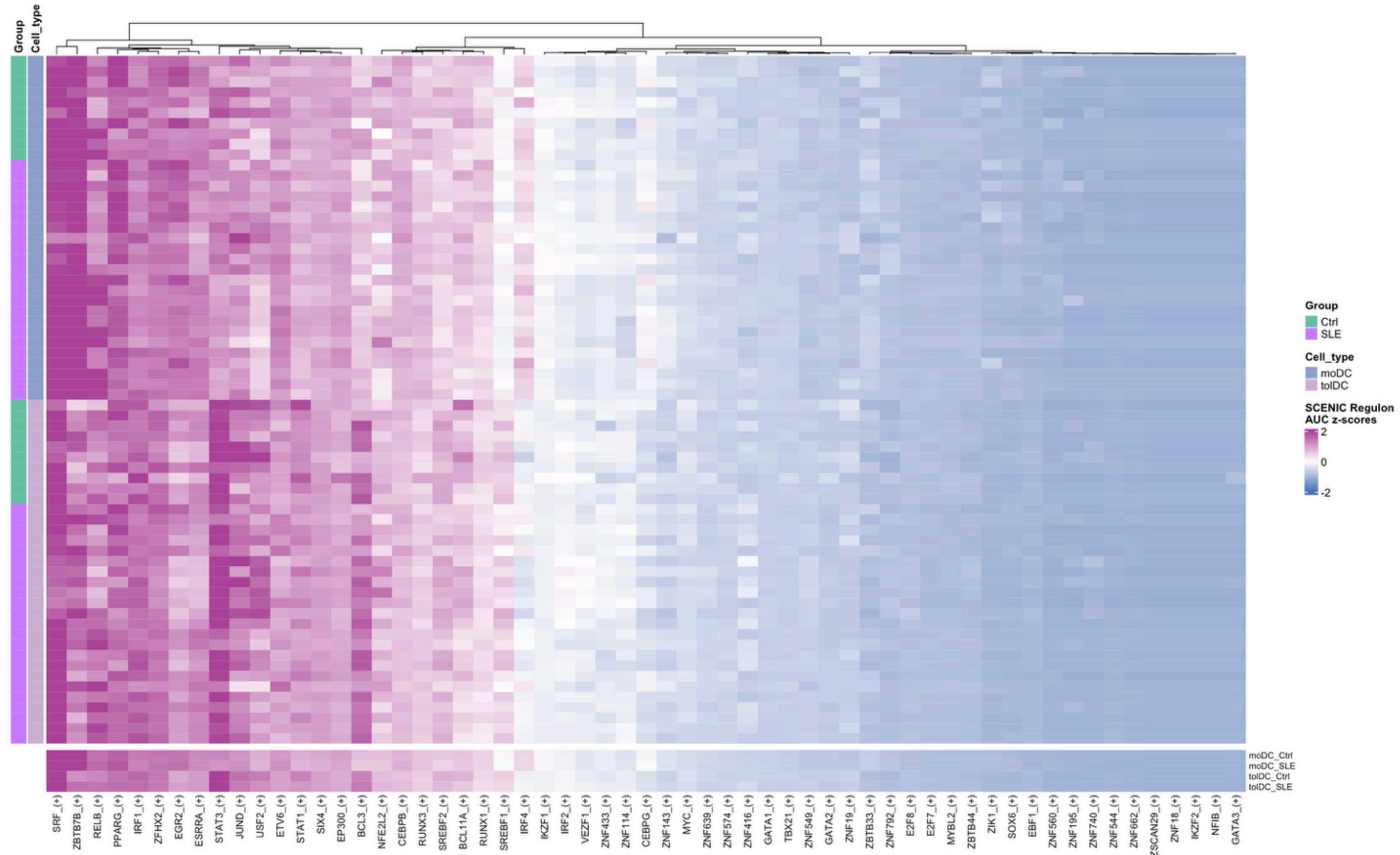


**Supplementary Figure S10. Regulon activity restricted to DEGs of interest (shared DEG from Figure 6A).** Activity of regulons that contain at least one differentially expressed gene (DEG) of interest among their targets in moDC and tolDC from SLE and control samples. For each regulon, we show the distribution of AUC z-scores across conditions.

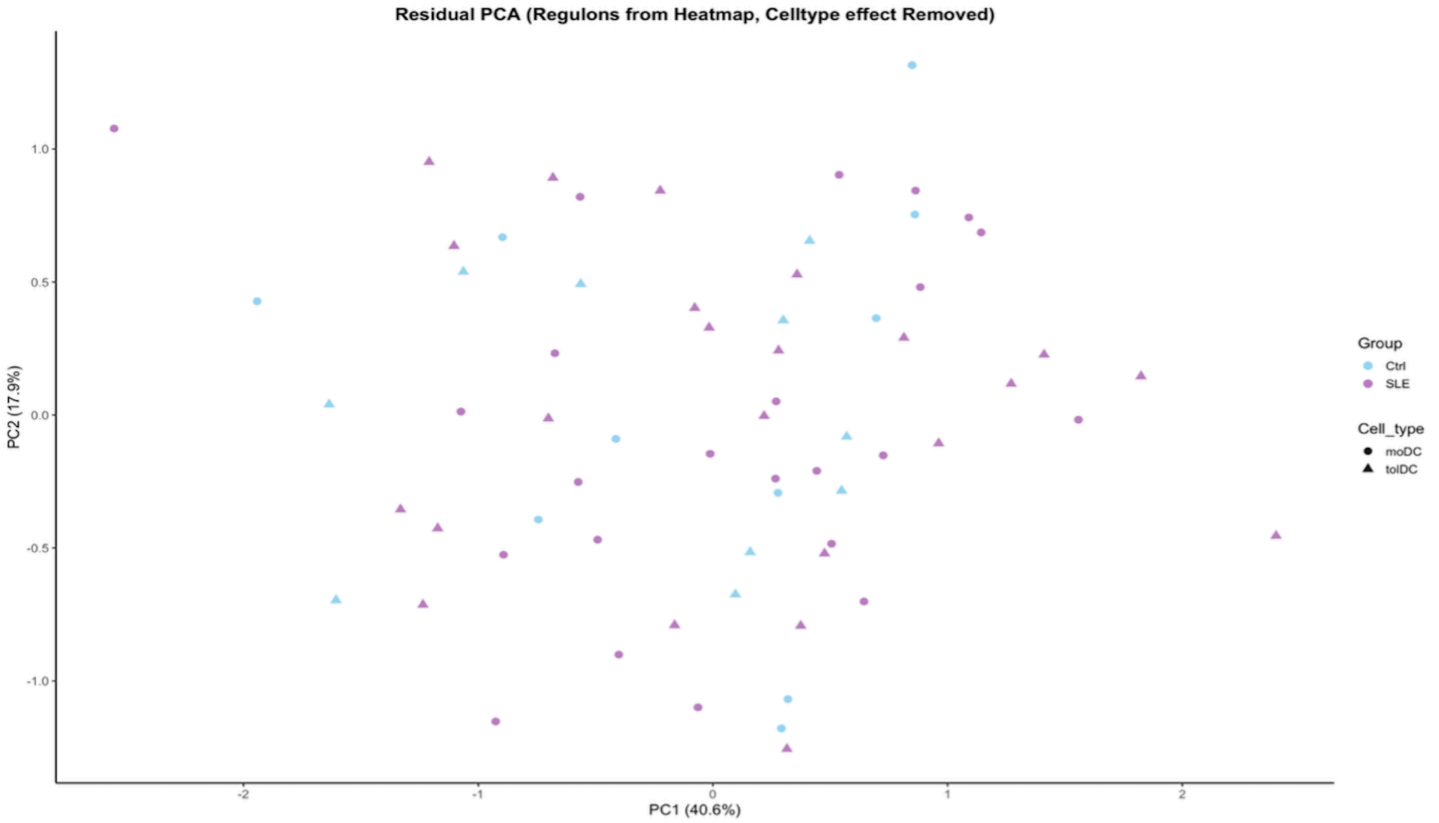


**Supplementary Figure S11. Residual PCA of regulon activity after removing cell-type effects**
Residual principal component analysis (PCA) of regulon activity after regressing out the effect of cell type from the AUC matrix using a linear model. In the original PCA using the full set of DEGs, the main axis of variation was driven almost exclusively by cell identity, with no clear segregation between SLE and control samples. The residual PCA, computed on regulon AUC scores after removing the cell-type effect, similarly failed to reveal a distinct separation by disease status. These results indicate that the dominant structure of the regulon landscape is shaped by cell differentiation state (moDC vs tolDC), and that any disease-associated differences occur as more subtle shifts in specific regulons rather than as a global reorganization of regulatory programs.

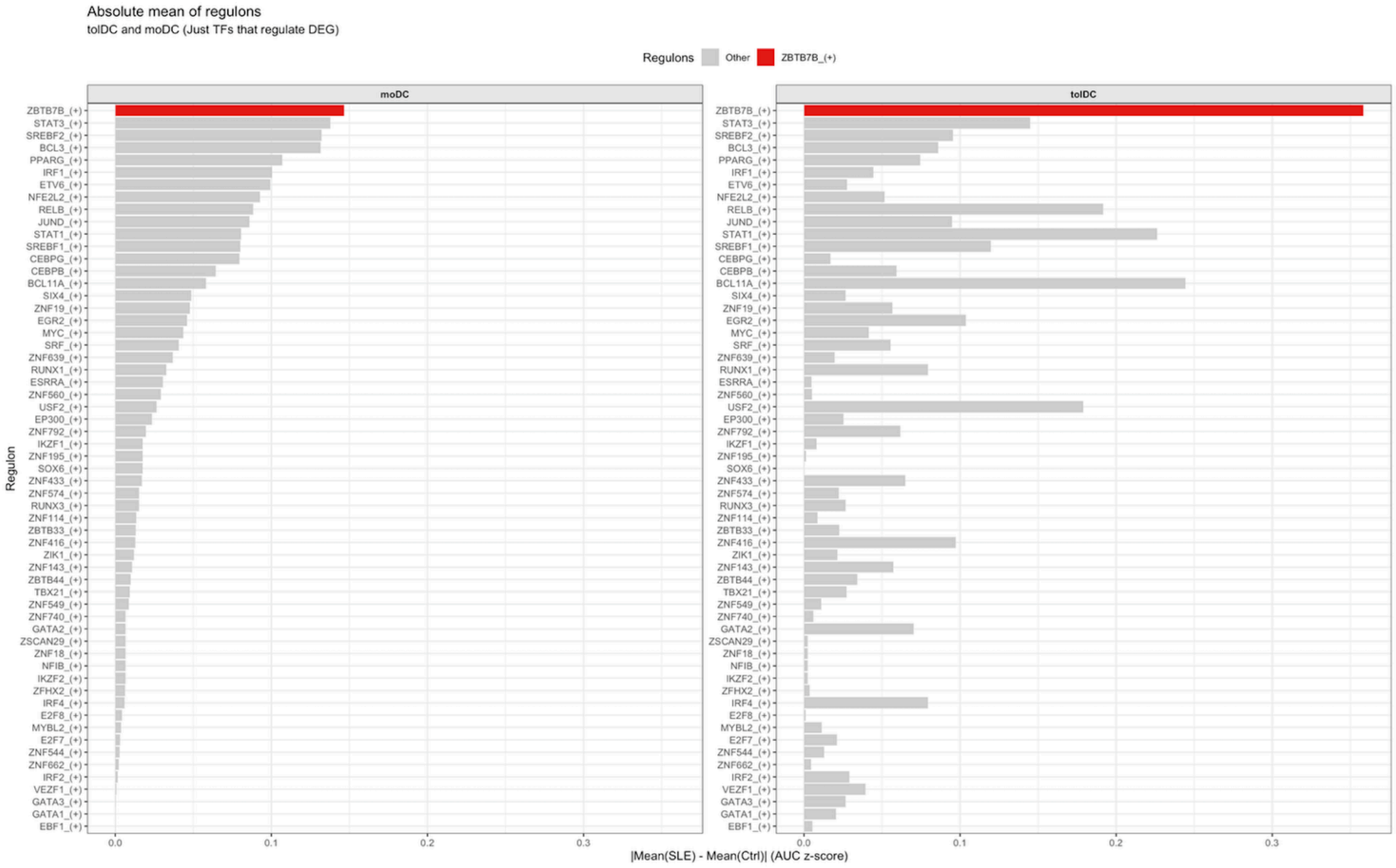


**Supplementary Figure S12. Absolute difference in regulon activity between SLE and controls**

For each regulon, the absolute difference in mean AUC z-score between SLE and control samples, computed separately for moDC and tolDC. This metric captures the magnitude of condition-specific changes in regulon activity while ignoring directionality, thereby allowing the identification of regulons that are most strongly perturbed in SLE regardless of whether they are up- or down-regulated. Within this framework, ZBTB7B emerges as the regulon with the largest difference between groups, particularly pronounced in tolDC, which led us to prioritize this transcription factor for downstream analyses.
